# Exploring the genetic landscape of ciprofloxacin-induced DNA supercompaction in *Escherichia coli*

**DOI:** 10.1101/2025.07.12.663469

**Authors:** Krister Vikedal, Natalia Berges, Ida Mathilde Marstein Riisnæs, Synnøve Brandt Ræder, Jørgen Vildershøj Bjørnholt, Magnar Bjørås, Kirsten Skarstad, Emily Helgesen, James Alexander Booth

## Abstract

DNA-damaging antibiotics like ciprofloxacin induce extensive double-strand breaks in *Escherichia coli*, triggering the SOS response and leading to DNA supercompaction. To uncover genes involved in this process beyond the previously identified core factors of *recN* and *recA*, we conducted a genome-wide screening using high-content imaging and machine learning-assisted image classification on nearly 4,000 *E. coli* strains, including the Keio collection’s single-gene deletion strains and additional in-house strains. Our investigation revealed novel genes contributing to supercompaction, with effects varying by genetic background. DNA supercompaction was consistently observed across eight clinical isolates from diverse bacterial species, underscoring the conservation of this cellular response. Our findings confirm RecN and RecA as primary drivers of DNA supercompaction. Additionally, we identified repair genes and novel genes that contribute to the response, especially in clinical *E. coli* strains. Notably, select hit gene deletions, including those for the membrane-associated proteins Hfq and YaiW, reduced RecN colocalization with the nucleoid, indicating a potential mechanism by which these genes impair supercompaction. Altogether, this work demonstrates that high-content imaging combined with automated analysis provides a powerful approach to explore population-level nucleoid dynamics and DNA damage responses, opening new avenues for understanding cellular processes and combating bacterial infections.

**GRAPHICAL ABSTRACT:** 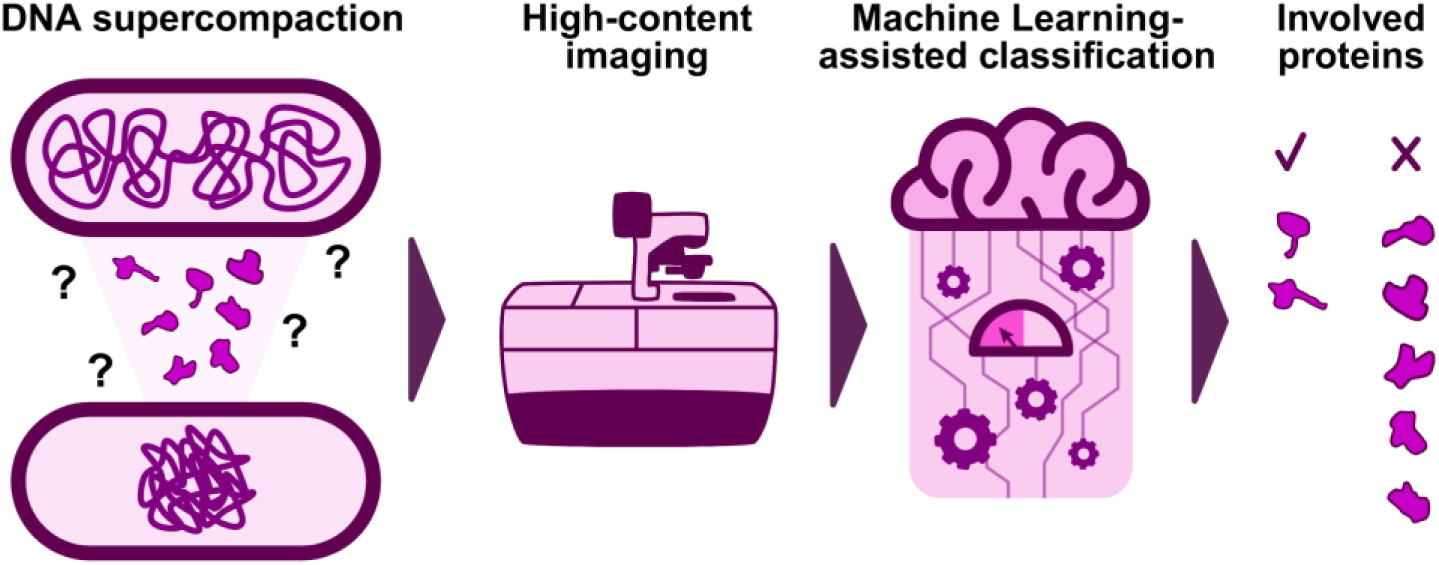

## INTRODUCTION

The emergence and spread of fluoroquinolone antibiotic resistance represents a critical challenge to global health (1). Ciprofloxacin (CIP), a widely prescribed fluoroquinolone, damages DNA by forming adducts with DNA gyrase and Topoisomerase IV, thereby blocking the progression of the replication machinery (2–4). These effects ultimately result in the formation of DNA double-strand breaks (DSBs) and trigger the major bacterial DNA damage response known as the SOS response (2). When there is no DNA damage, the transcription factor LexA binds to regions upstream of DNA repair genes, thereby repressing their transcription. Upon DNA damage, RecA binds to single-stranded DNA, where it not only participates in DNA repair but also triggers the SOS response by promoting the autocleavage of the LexA repressor (5, 6).

The SOS response constitutes a sophisticated cellular program that coordinates multiple survival mechanisms (7). Upon activation, this system triggers extensive changes in gene expression, leading to the production of DNA repair proteins, protective factors, and a multitude of uncharacterized gene products (8, 9). For instance, the SOS response induces the expression of proteins necessary for complex DSB repair mechanisms centered around homologous recombination (10). This process begins when the RecBCD complex processes DNA ends at DSBs, generating single-stranded DNA that serves as a substrate for RecA protein loading (11). RecA then catalyzes the search for and pairing with intact homologous sequences, enabling repair of the damaged DNA region (12, 13).

The SMC-like protein RecN has also been established as an important factor in DSB repair (14, 15) and stands out as one of the most highly expressed proteins during the SOS response (8, 9). Its protein levels are tightly controlled through rapid degradation by the ClpXP protease system (16, 17). Although RecN-deficient cells show increased sensitivity to CIP-induced DNA damage (18), the precise mechanism by which RecN operates and contributes to DSB repair remains incompletely understood.

A striking aspect of the bacterial response to DNA damage is nucleoid compaction, observed following exposure to various genotoxic agents including UV irradiation, mitomycin C, bleomycin, and fluoroquinolones such as CIP and ofloxacin (18–22). Recently, we discovered that severe DNA damage induced by agents like CIP triggers a stepwise reorganization of the nucleoid, a process we termed “DNA supercompaction” (18). Upon severe DNA damage, the cell’s nucleoid rapidly transitions from its multifocal distribution— common in unchallenged conditions—through quarter-position compaction, culminating in a dense, persistent midcell compaction. Quarter-position compaction involves the formation of two distinct nucleoid lobes situated in separate cell halves. Although the DNA supercompaction process depends on both RecA and RecN, a quarter-position compaction occurs even in their absence, albeit in a temporally dysregulated manner (18). The independence from RecA and RecN suggests a likely involvement of genetic factors that are not yet identified in the DNA supercompaction process.

The Keio collection of *Escherichia coli* strains, featuring single-gene deletions of all nonessential genes from the BW25113 background (23–25), has been widely used to explore various bacterial phenomena (26–31). In this study, we systematically screened the Keio collection and additional in-house strains to pinpoint genetic factors important for DNA supercompaction following CIP exposure. Utilizing high-content imaging and machine learning-assisted image analysis, we identified candidate strains with compaction phenotypes like either *ΔrecN* or unchallenged strains. Following a stringent validation process, we identified 15 hit strains, featuring deletions of both known DNA repair genes and novel genes unassociated with DNA compaction or repair, such as *ΔdusB*, *Δhfq*, *ΔyaiW* and *ΔydeE*. Notably, these deletions exhibited varying effects on compaction progression across different genetic backgrounds, with the clinical reference background CCUG17620 consistently displaying impaired supercompaction for all examined deletions. Additionally, several hit gene deletions appeared to affect RecN dynamics in the supercompaction process. These findings indicate that, while RecN and RecA are the principal facilitators of DNA supercompaction, a broader network of genetic factors are involved in nucleoid reorganization in response to DNA damage.

## MATERIALS AND METHODS

### Strains used in screening and growth conditions

For the screening, we included a total of 3862 *E. coli* strains, comprising 3797 non-essential single-gene deletion strains from the Keio collection (23) and 65 strains from our in-house library (Supplementary Table S1). The in-house strains, featuring various genetic backgrounds with single or multiple gene mutations and deletions, were included due to their potential relevance to the DNA supercompaction process. To facilitate high-throughput handling, strains were distributed across a series of 13 different 384-well plates (Merck Life Science no. P5991), excluding outer wells. Each plate contained two samples of a wild-type strain (BW25113) (32) and a *ΔrecN* strain (JW5416) (23) as controls for the DNA supercompaction process (18).

Strains were cultured in LB medium with relevant antibiotics (30 µL per well) within 384-well plates and incubated at 37°C in sealed bags with high humidity to prevent drying. Kanamycin (30 µg/mL) was used for Keio collection strains, while for the in-house strains, antibiotics as specified in Supplementary Table S1 were used at the following concentrations: 20 µg/mL Chloramphenicol, 5 µg/mL Tetracycline, and 50 µg/mL Ampicillin.

### Sample preparation for screening

Overnight cultures (ONCs) were inoculated directly from freeze stock using a 96-pin Multi-Blot Replicator (V & P Scientific no. VP 408FP6 and VP381). On the day of imaging, new cultures were inoculated from the ONCs and incubated until the OD_600_ of control samples reached 0.4. Next, CIP (10 µg/mL) was added to all wells, excluding one sample for each of the control strains, to induce severe DNA damage and trigger a rapid, ordered DNA supercompaction response (18). Samples were incubated with CIP for 15-20 minutes before fixation with ethanol (50% final concentration).

For imaging, glass-bottomed 384-well plates (Fisher scientific no. 10687354) were pre-treated with 25 µL Poly-L-Lysine (Merck Life Science no. P8920) for 8 minutes and washed once with PBS. Fixed samples (15 µL) were transferred to these plates and DNA was stained with 5 µg/mL Hoechst 33258. During staining, plates were centrifuged at 700 x *g* for 10 minutes to ensure sufficient cell adhesion to the bottom of the well. After centrifugation, the supernatant was carefully removed, wells were washed once with PBS, before finally adding 25 µL PBS. Plates with samples were brought to the microscope for imaging immediately after preparation and maintained at room temperature throughout the procedure.

### High-content imaging for screening

Samples in glass-bottomed 384-well plates were imaged using an ImageXpress Micro Confocal High-Content Imaging System (Molecular Devices), equipped with a 60x air objective (Nikon CFI Plan Apochromat Lambda 60XC, 0.95 NA) and an Andor Zyla sCMOS camera. Imaging and focusing were fully automatic, though each plate’s automatic focus setup was verified and adjusted as needed. Four locations per well were imaged, spaced evenly by 600 µm, using two channels with auto-shading correction: a transmitted light (TL) channel for detecting cell outlines and a fluorescence channel for Hoechst 33258, with excitation at 350-404 nm and emission collection at 417-477 nm. The optimal focus for each location was automatically selected from a Z-stack of three images separated by 0.4 µm in each direction from the default focus position. Edge wells were excluded from imaging due to interference between the objective and the plate skirt (according to microscope manual), a limitation accounted for during strain plate preparation.

### Image analysis and strain classification in screening

We utilized the open-source software CellProfiler (version 4, (33)) to preprocess and analyze images from the screening, together with its companion tool CellProfiler Analyst (version 3, (34)) for training machine learning models for cell classification. Full pipelines are available in our Zenodo repository (see Data Availability). All images, with a pixel size of 115 nm x 115 nm, were preprocessed to remove background noise. The TL channel was enhanced to isolate dark hole elements 7-35 pixels in diameter and suppressed for elements smaller than 6 pixels wide. The fluorescence channel underwent enhancement for speckle elements up to 18 pixels in diameter. Cells were segmented from these preprocessed TL channels by identifying objects with diameters of 8-35 pixels, using a global “Robust Background” thresholding method with mode averaging. Cell segmentation did not involve splitting of cell clusters. DNA foci were identified from corresponding fluorescence channels by detecting objects smaller than 35 pixels in diameter, employing a global “Robust Background” thresholding method with median averaging. Clustered DNA foci were distinguished by separating local intensity peaks at least 4 pixels apart after application of a smoothing filter. Holes in identified DNA objects were filled, and only cell objects collocating with DNA foci were included in subsequent analyses. A range of measurements were collected for the segmented cell objects and their DNA foci, encompassing features such as size, shape, and DNA distribution (see Supplementary Material).

Two machine learning models were developed for cell classification using CellProfiler Analyst: the Single-Cell model and the Phenotype model. These models were trained using a 15-parameter FastGentleBoosting classification strategy within CellProfiler Analyst, utilizing both simple and complex measurements from CellProfiler analyses (training details in Supplementary Material). The Single-Cell model refined cell segmentation accuracy by excluding clusters, imaging artefacts, and out-of-focus cells, establishing a solid basis for subsequent phenotype classifications. The Phenotype model classified cells by their DNA supercompaction phenotypes—defined in this study as wild-type, *ΔrecN*, and unchallenged. This classification provided insights into phenotype prevalence across samples and was fundamental for enrichment scoring and candidate strain identification.

To efficiently process the large volume of images generated in the screening, we employed CellProfiler’s batch processing capabilities on cluster computers to automate image analysis. This approach generated extensive spreadsheets with measurements for various features of every individual cell. We developed a Python script to compile these results from all cells within a given sample, conduct statistical analyses, and organize the data into a manageable format for further review. Enrichment scores for DNA supercompaction phenotypes were calculated for every sample (see Supplementary Material) using the same method employed by CellProfiler Analyst (35). All relevant scripts are available in our Zenodo repository (see Data Availability).

### Growth conditions for individual culturing

Strains were cultured individually to ensure they reached exponential growth prior to treatments and fixation. Initially, frozen stocks were streaked onto LB-agar plates and incubated overnight at 37°C. The following day, fresh growth medium was inoculated with 5-10 colonies from each strain and incubated overnight with shaking. At the start of an experiment, ONCs were diluted depending on cell density (1:500 or 1:200 for most) in fresh growth medium and cultured to exponential growth (OD_600_ around 0.2-0.4) before applying treatments. LB medium was generally used for culturing, except for in a screening validation experiment where AB minimal medium (36) supplemented with 0.5% casamino acids, 0.2% glucose, and 1 µg/mL thiamine was utilized to provide a different growth condition (37).

Cells were exposed to CIP at a dose of 10 µg/mL for 20 minutes, unless otherwise specified, to effectively differentiate the compaction phenotypes among strains (18). Ethanol at a 50% final concentration was used for cell fixation. Keio collection strains were cultured with 30 µg/mL Kanamycin, while in-house and constructed strains were cultured using antibiotics as indicated in Table 1 and Supplementary Table S1, at the following concentrations: 100 µg/mL Ampicillin, 20 µg/mL Chloramphenicol, 30 µg/mL Kanamycin and 5 µg/mL Tetracycline.

**Table 1.**
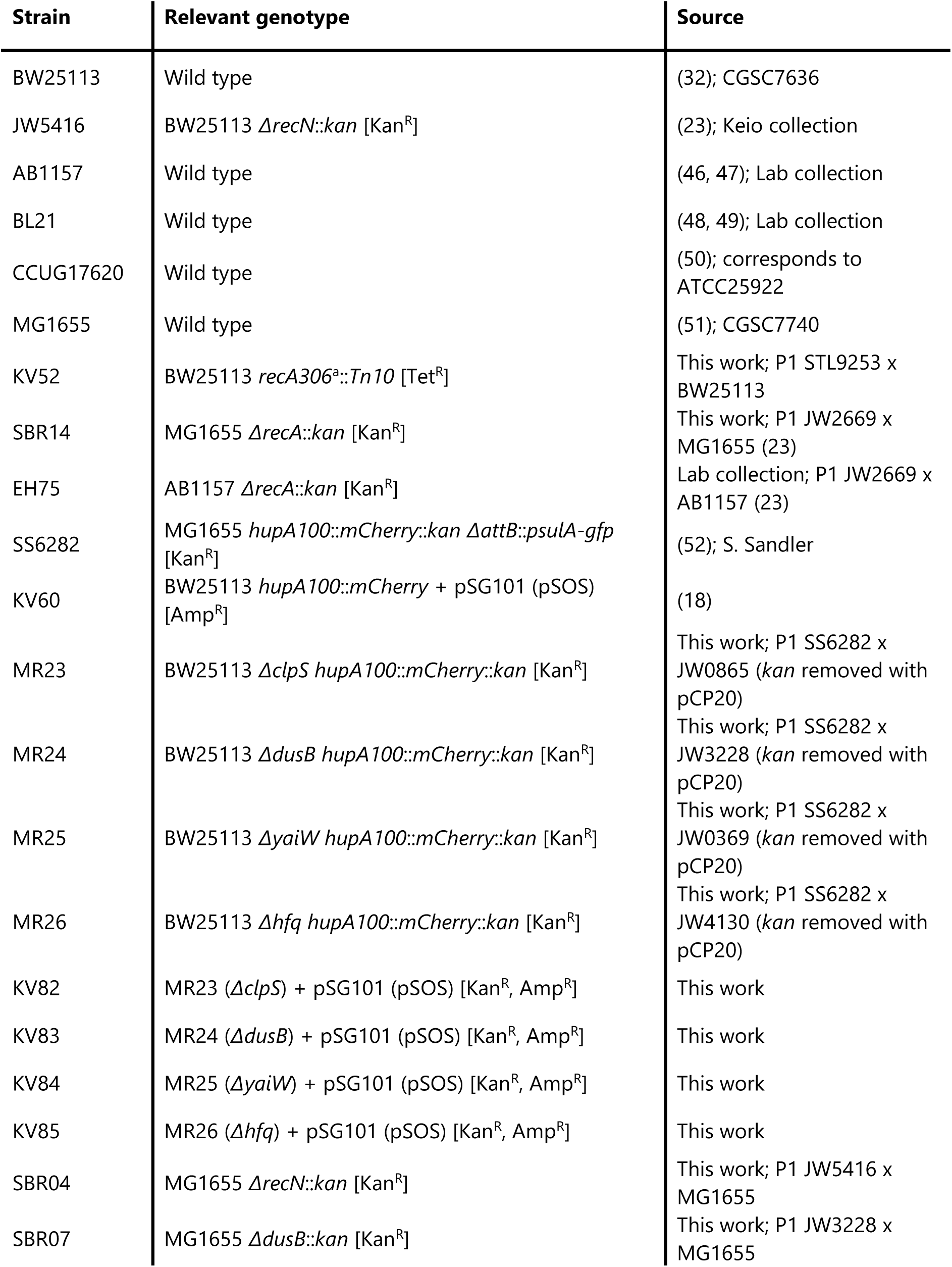
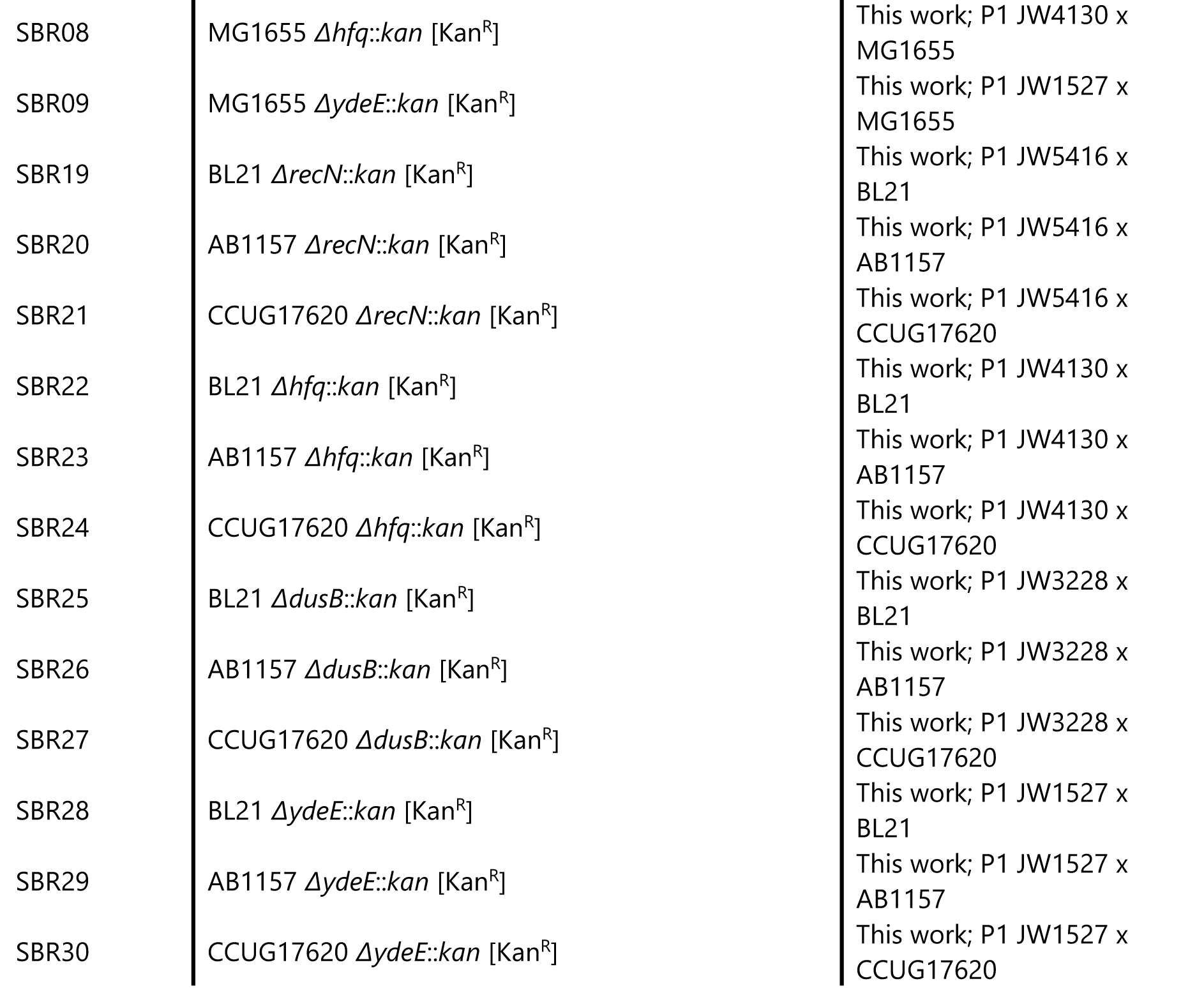
Strains used in this study to investigate the role of hit strain mutations and deletions in DNA supercompaction. P1 transduction is shown as: P1 Donor x Recipient. Relevant antibiotic resistances are shown in brackets: Amp^R^, Ampicillin resistance; Cam^R^, Chloramphenicol resistance; Kan^R^, Kanamycin resistance; Tet^R^, Tetracycline resistance. ^a^ The *recA306* notation refers to the *Δ(srl-recA)306* mutation, which only leaves a small portion of the *recA* gene and can be considered a deletion of *recA* **(43–45)**.

### Fluorescence microscopy for validation of screening candidate strains

Fixed samples of individually cultured strains were imaged with fluorescence microscopy to assess DNA compaction phenotypes following CIP exposure, ensuring controlled conditions for validation of candidate strains. This approach utilized a Leica DM6000 B microscope equipped with a 100x oil objective (Leica HCX Plan Apochromat 100x 1.40 NA, PH3 CS), a Leica EL6000 metal-halide light source, and a Hamamatsu C9100-14 EM-CCD camera. After treatment and fixation, cells were stained with 5 µg/mL Hoechst 33258 in PBS for 10 minutes, washed once with PBS, and concentrated 2-4 times in a PBS resuspension. The stained cells (10 µL) were evenly distributed onto an agar pad pre-made within a Gene Frame (Thermo Scientific no. AB0576) on a glass slide and sealed with a coverslip after drying. Images were captured from 4-6 distinct locations across the agar pad to achieve comprehensive cell representation. Each image included a phase-contrast channel for cell outlines and a fluorescence channel for Hoechst 33258, with excitation at 334-406 nm and emission collection at 407-497 nm.

### Strain construction for investigation of DNA supercompaction

Strains constructed from screening hit strains to investigate the role of their mutations and deletions in DNA supercompaction are listed in Table 1. Standard P1 transduction procedures (38) were employed to transfer genetic mutations and deletions from these hit strains into various background strains, as well as to introduce *hupA100*::mCherry (39) chromosomally for DNA visualization during live-cell imaging. Electroporation transformation, as detailed in our previous study (18), was used to introduce plasmids required for studying RecN localization and dynamics, and for the SOS response assay. The pSG101 (pSOS) plasmid encoded GFP-RecN regulated by the native RecN promoter (17), while the p*lexA*-*gfp* reporter plasmid encoded GFP regulated by the *lexA* promoter (40, 41). In cases where both donor and recipient strains carried kanamycin resistance, the donor *kan*-cassette was removed using FLP recombinase (pCP20) (42).

### Live-cell spinning disk microscopy

To explore DNA supercompaction progression and RecN dynamics in selected hit strains, live-cell spinning disk microscopy was employed with the same methodology outlined in (18). In brief, the setup involved a Nikon Eclipse Ti2-E inverted microscope configured with a 60x oil objective, a CrestOptics X-Light V2 spinning disk confocal module (50:400 µm spinning disk), and a Lumencor Celeste multi-line laser, supplemented with two Teledyne Photometrics Kinetix sCMOS cameras. Temperature control was maintained through a stage-top incubator chamber and an objective heater. Strains were individually cultured to exponential phase, then exposed to CIP (10 µg/mL) and incubated for an additional minute. Immediately after, they were transferred onto LB½-agar pads (LB diluted 1:1 with Milli-Q water, with 1% agarose) on microscope slides within a Gene Frame (Thermo Scientific, no. AB0576), sealed with a cover slip (#1.5 thickness). Imaging commenced 10 minutes after CIP exposure, acquiring images from three location at 2-minute intervals while maintaining samples at 37°C. Fluorescence imaging employed three channels: a transmitted light channel for cell outlines, and GFP and mCherry fluorescence channels. GFP was excited at 477 nm with emission collected between 501-521 nm, while mCherry was excited at 546 nm with emission collected between 580-610 nm.

### Image analyses to characterize DNA distributions and RecN dynamics

Image processing and analyses followed protocols we established in our previous study (18). We utilized the open-source software Fiji (ImageJ) (53), alongside the plugins Coli-Inspector (54) and MicrobeJ (55), to assess DNA supercompaction and RecN dynamics.

The ObjectJ-based Coli-Inspector plugin (version 03f) (54) analyzed images to characterize DNA distribution through DNA profiles. Briefly, images were preprocessed to remove background noise, followed by cell segmentation with adjusted parameters for cell area, width, and circularity. CIP-exposed cells were grouped according to exposure duration. DNA profiles were determined by measuring average symmetrical fluorescence intensity along cells’ long axes, with DNA profile widths identified as the outer bounds of the fluorescence peaks at 80% of their maximum intensity. These measurements were made and plotted with GraphPad Prism (version 10.4.1), to visualize the DNA distribution along the cells’ long axes.

MicrobeJ (beta-version 5.13p (20)) (55) facilitated the creation of kymograph heat maps and the quantification of RecN foci and their colocalization with HU-mCherry foci. In brief, preprocessing removed images’ background noise, followed by cell segmentation using the “Medial Axis” method, with limitations applied to cell area, length, width, curvature, and sinuosity. Kymograph heat maps, generated using the ShapePlot tool, summarized the average fluorescence distribution within cells. GFP-RecN foci were detected within segmented cells for quantification, with their positions compared to HU-mCherry foci locations to assess RecN colocalization with DNA. The resulting data were transferred to GraphPad Prism (version 10.4.1) for plotting.

### SOS response assay

To assess SOS response activity in screening hit strains, the SOS response assay involved transformation of these strains with a p*lexA*-*gfp* reporter plasmid, which conferred either chloramphenicol or kanamycin resistance depending on the strains’ existing resistance genes. The plasmid conveys GFP fluorescence, regulated by the *lexA* promoter, was quantified using flow cytometry. Strains were cultured to exponential growth in LB medium. Samples of 6 µL were collected every 30 minutes, from just before until 120 minutes after CIP exposure, when the wild-type control sample’s SOS response activity plateaued. These samples were diluted in 200 µL PBS within wells of a 96-well plate, adjusting the sampled cell culture volume if necessary to achieve approximately 2,000 events per µL. The plate with diluted samples was immediately brought to an Accuri C6 flow cytometer (BD Bioscience) for measurement of integrated GFP fluorescence signal from 20,000 cells per sample. GFP was excited at 488 nm and emission was collected at 515-545 nm. A detection threshold for forward scatter height was applied to exclude smaller particles from analyses. SOS response activity was quantified by normalizing the sample’s mean GFP fluorescence to the mean forward scatter area (mean cell size), under the assumption that larger cells contain more GFP.

### UV exposure of hit strains

Hit strains were irradiated with UV to assess DNA supercompaction and deletion-dependent survival effects. Each strain’s ONC was diluted to achieve new cultures with an OD_600_ of 0.01 in fresh LB medium containing appropriate antibiotics. These cultures were incubated at 37°C until reaching exponential growth at an OD_600_ around 0.2. Cultures were then split into two subcultures: one remained unchallenged, while the other was UV irradiated. Both subcultures were pelleted by centrifugation at 20,000 x *g* for 30 seconds at 37°C and resuspended in PBS to prevent UV absorption by the LB medium. Suspensions were gently pipetted onto the center of 90 mm-diameter Petri dishes. The exposed subculture received a UV dose of 5 J/m^2^ at 254 nm using a UV crosslinker (UVP CL-1000). Following UV exposure, subcultures were collected, pelleted by centrifugation, and resuspended in LB medium with appropriate antibiotics. For survival assays, 100 µL of each resuspension underwent serial ten-fold dilution in LB medium within 96-well plates. For imaging, remaining cell suspensions were incubated in a water bath at 37°C with shaking for 15 minutes before fixation with an equal volume of ethanol. These samples were subsequently imaged using the same fluorescence microscopy approach as employed to validate screening candidate strains.

### Survival assays

Survival assays examined the tolerance of hit strain samples to CIP or UV exposure, relative to unchallenged samples. Strains were cultured until their OD_600_ reached 0.2, at which point cultures were split into two subcultures: one remained unchallenged, while the other was exposed to either CIP or UV. For CIP exposure, a 10 µg/mL dose was employed to assay the time-dependent survival of strains, as detailed in our previous study (18). Samples were collected from each subculture prior to treatment and at time points ranging from 1 to 60 minutes after exposure. For UV exposure, wild-type (BW25113) and *ΔrecN* (JW5416) strains were assayed in a dose-dependent manner using UV doses ranging from 5 to 70 J/m^2^, applying the exposure method described earlier. The survival of hit strains was only assayed at a single dose of 5 J/m^2^.

Ten-fold serial dilutions of both exposed and unchallenged subcultures were prepared, with each dilution spotted onto LB-agar plates containing relevant antibiotics for selection. Plates were incubated overnight at 37°C. For each strain, colonies were counted at the lowest dilution where distinguishable colonies appeared, and colony forming units (CFU) per mL were calculated for both subcultures. Relative survival after treatment was expressed as the ratio of CFU/mL between exposed and unchallenged subcultures.

### Statistical analysis

The initial screening was performed in three biological replicates, complemented by re-imaging candidate strains and follow-up experiments to ensure a sufficient number of replicates supported qualified decision-making when selecting hit strains. Details regarding testing of machine learning models and calculation of enrichment scores for screening candidates are provided in the Supplementary Material. The number of biological replicates for the experiments investigating the role of hit strain mutations and deletions in DNA supercompaction is indicated in the corresponding figure captions. An ordinary one-way ANOVA with Dunnett correction for multiple comparisons was used to analyze differences in SOS response activity, as well as differences in DNA profile widths and survival rates following 5 J/m^2^ UV irradiation. These analyses were conducted using GraphPad Prism (version 10.4.1). Figures employ color-coding to visually indicate significant differences between tested strains, while asterisks denote the level of statistical significance. Time- and dose-dependent survival data after CIP exposure and UV irradiation, respectively, were analyzed in R (version 4.4.1). Survival values, relative to unchallenged parallels, were log_10_- transformed and fitted by linear mixed-effects models including only main effects, with random intercepts for biological replicates (see Zenodo repository in Data Availability). Estimates with 95% confidence intervals are presented for fixed effects. Two-tailed *P*-values below 0.05 were considered significant and indicated with asterisks as follows: \**P* ≤ 0.05; \*\**P* ≤ 0.01; \*\*\**P* ≤ 0.001; \*\*\*\**P* ≤ 0.0001.

## RESULTS

### Machine learning-assisted screening to identify *E. coli* strains with impaired DNA supercompaction progression

To identify genes involved in DNA supercompaction following CIP exposure, we screened 3797 single-gene deletion *E. coli* strains from the Keio collection (23), along with 65 additional in-house strains (Supplementary Table S1; overview of screening procedure in Fig. 1). The strains were cultured to exponential phase in 384-well plates and then exposed to 10 µg/mL CIP for 15-20 minutes, followed by ethanol fixation, Hoechst 33258 DNA staining, and high-content imaging. Our previous work demonstrated that CIP exposure triggers DNA supercompaction in wild-type *E. coli* cells (BW25113)—a response characterized by the nucleoid’s morphology transitioning from a multifocal distribution under unchallenged conditions, through quarter-position compaction, to a dense midcell compaction (18). In contrast, *ΔrecN* cells fail to progress through this response, instead exhibiting a consistent form of quarter-position compaction (18). To accurately assess phenotypic responses of the screened strains, we included three controls for comparison: wild-type cells exposed to CIP, *ΔrecN* cells exposed to CIP, and unchallenged samples of wild-type and *ΔrecN* cells.

**Figure 1.**
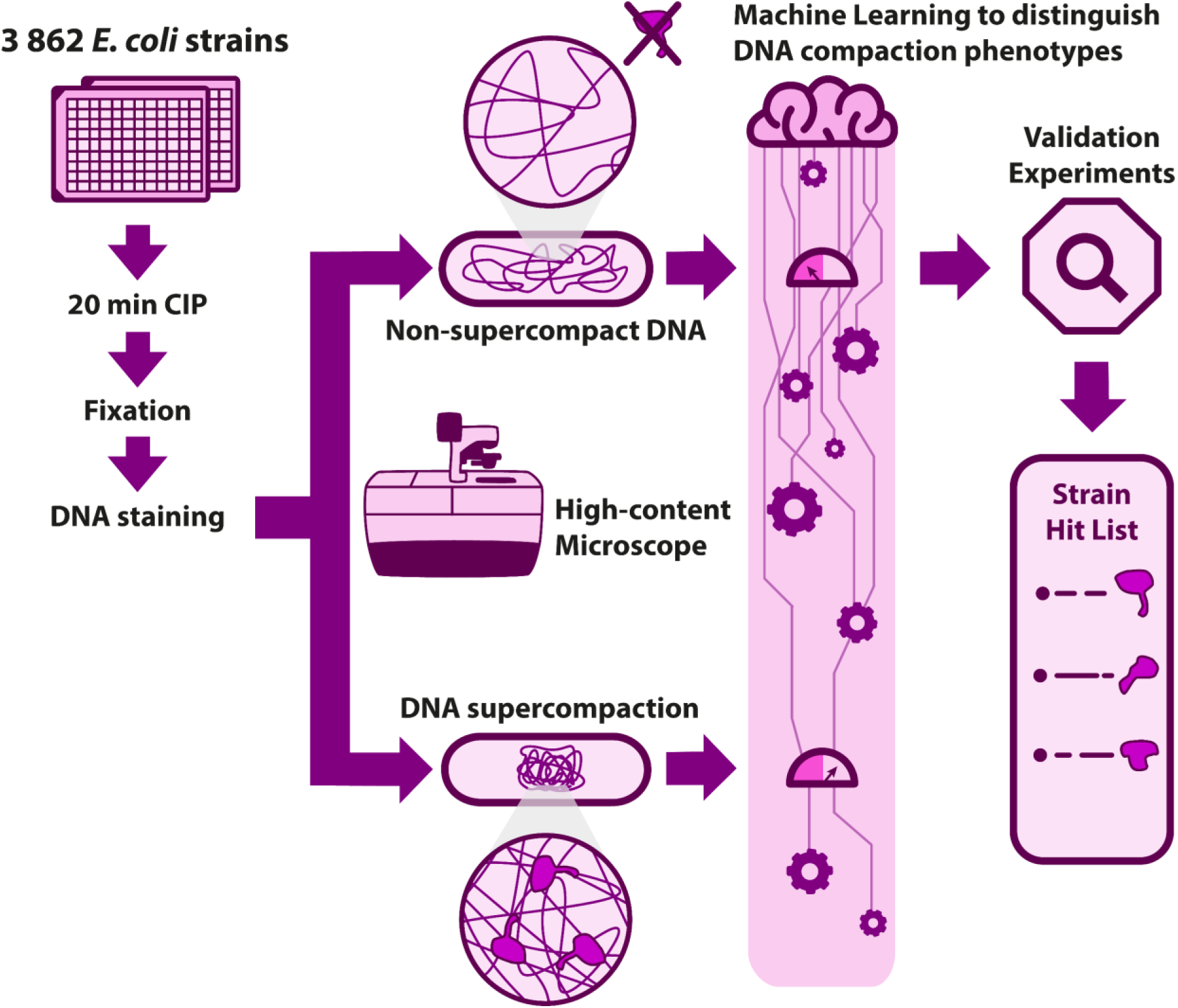
Overview of screening procedure used to identify gene deletions affecting the DNA supercompaction response in *E. coli*. We cultured strains, along with controls, to exponential phase in 384-well plates. Cells were then exposed to CIP (10 µg/mL) for 20 minutes, fixed with ethanol, and stained with Hoechst 33258 (5 µg/mL) to visualize DNA. These stained samples were then transferred to glass-bottomed 384-well plates for automated, high-content imaging. A machine-learning assisted CellProfiler pipeline performed cellular measurements and classified their compaction phenotype. Strains showing impaired DNA supercompaction (*ΔrecN* or unchallenged compaction phenotypes) were identified as candidates for further validation experiments. Following these experiments, we conducted subjective evaluations of the candidates, ultimately selecting those with clearly impaired DNA supercompaction as hit strains.

High-content imaging produced nearly 50,000 images, which were analyzed in a high-throughput manner using CellProfiler (version 4, (33)) with batch processing on a cluster computer. Initially, we attempted to classify control sample phenotypes—wild-type, *ΔrecN*, and unchallenged (Fig. 2A)—using simple measurements like DNA foci count and relative area, in line with our previous work (18). However, we did not achieve optimal focus for each sample with the automated imaging setup of the screening, rendering these simple measurements insufficient for consistent classification. To overcome this obstacle, we employed machine learning models to distinguish compaction phenotypes more effectively.

**Figure 2.**
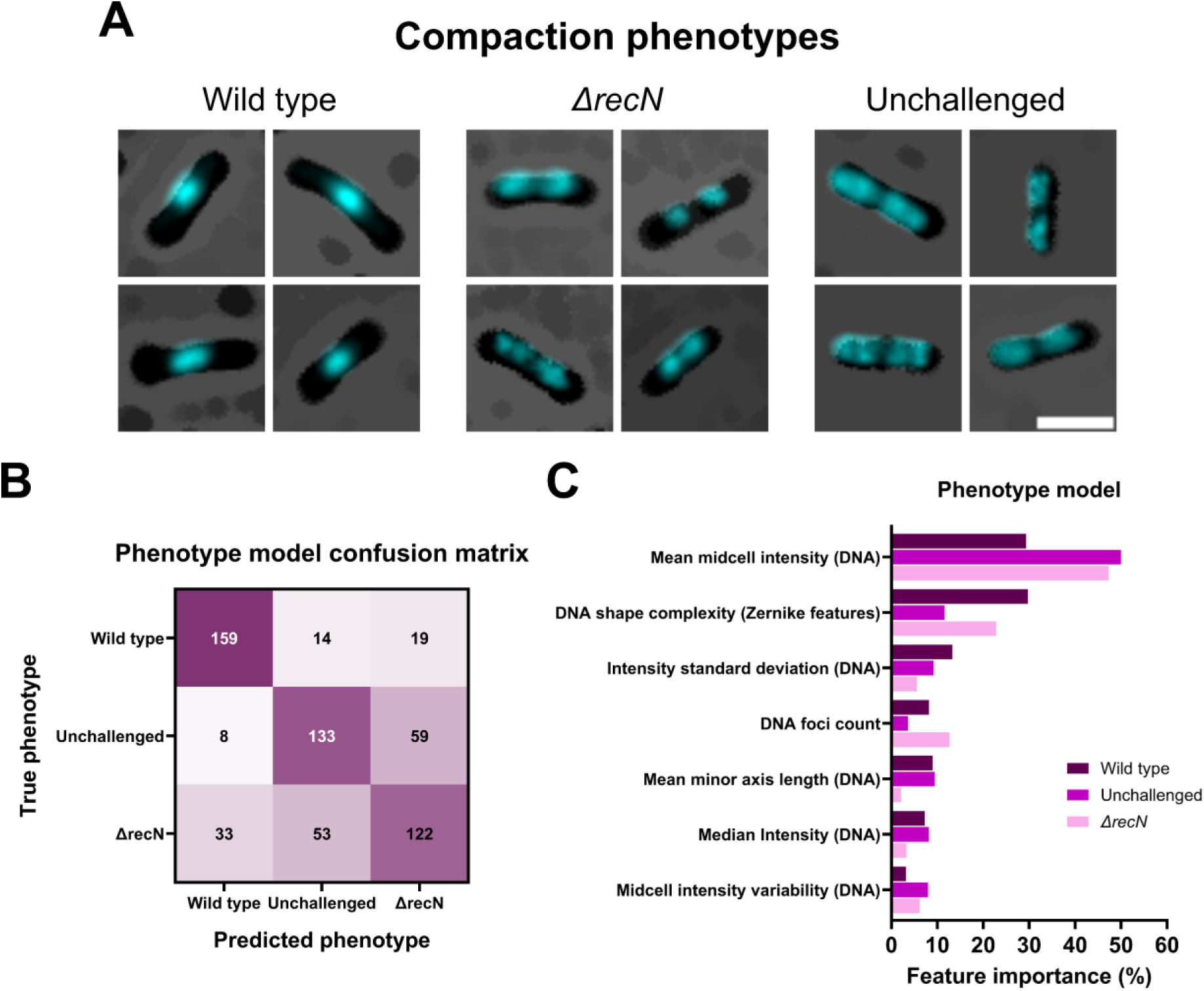
The Phenotype model was trained to differentiate the three DNA compaction phenotypes found in control samples—wild-type, *ΔrecN*, and unchallenged. (**A**) Images of representative cells from those samples, preprocessed according to the CellProfiler pipeline. Wild-type cells exhibit midcell compaction, the end-point of DNA supercompaction. The *ΔrecN* cells display quarter-position compaction, while unchallenged cells show multifocal distribution, both indicating impaired DNA supercompaction for CIP-exposed cells. Scale bar is 3 µm. (**B**) Confusion matrix for the Phenotype model quantifying its prediction performance. The matrix displays the true phenotypes for 200 cells per model-predicted phenotype. (**C**) Feature importance quantification for the Phenotype model, illustrating contribution of various parameters to classification of each compaction phenotype. Parameters are grouped based on similarity to emphasize which cellular features are critical for accurate phenotype differentiation.

We trained two machine learning models on 2,000 objects (cells) each from a subset of control sample images: a Single-Cell model to identify single cells for further analysis and exclude irrelevant objects, and a Phenotype model for classifying these cells’ compaction phenotype (training details in Supplementary Material). The FastGentleBoosting classification strategy of CellProfiler Analyst (version 3, (34)) provided reproducible and transparent models, facilitating easy interpretation of features and measurements used for classification. The Single-Cell model effectively excluded irrelevant objects, achieving an F_1_ score of 0.84, with cellular elongation as the key feature for classification (Supplementary Fig. S1 and Table S2). The Phenotype model reliably classified the wild-type phenotype with an F_1_ score of 0.81, though differentiation between *ΔrecN* and unchallenged phenotypes was less accurate (Fig. 2B; for detailed testing results see Supplementary Material). Nonetheless, the model’s performance was satisfactory since strains with impaired DNA supercompaction are expected to exhibit either the *ΔrecN* or unchallenged phenotypes. For this model, the mean midcell Hoechst 33258 intensity was the most important feature (Fig. 2C and Supplementary Table S3; feature importance calculation is detailed in Supplementary Material, see also Zenodo repository in Data Availability). The model also incorporated multiple Zernike features among its parameters (56, 57)—measurements of DNA shape complexity—that played significant roles in differentiating the DNA distribution across phenotypes. While DNA foci count contributed to classification accuracy, it represented only about 10% of feature importance (Fig. 2C), underscoring the need to combine various simple and complex measurements for precise classification.

To assess the practical performance of the machine learning-assisted classification, we applied the two models to classify the compaction phenotypes of 83 control samples— half of those included in the screening. The Phenotype model classified every individual cell, enabling sample-level phenotype classification through the calculation of enrichments scores (see Supplementary Material for details). Overall, 63 samples (75.9%) were correctly classified, while only two samples (2.4%) were misclassified, and 18 samples (21.7%) remained unclassified. Notably, all except one of the CIP-exposed wild-type controls were correctly classified with the wild-type phenotype. We were satisfied with the performance of the models and integrated them into our CellProfiler pipeline for batch processing of all strain images from the screening (details in Supplementary Material).

Upon screening completion, we identified 156 candidate strains exhibiting impaired DNA supercompaction progression following CIP exposure (Supplementary Table S4; detailed selection criteria in Supplementary Material). Additionally, we experienced insufficient growth for some strains, which led to subsequent challenges with phenotype assessment. This observation aligns with previous reports on some Keio collection strains growing slower than the wild type (28, 30, 58). As outlined in the Supplementary Material, 93 strains displayed poor growth in our screening (Supplementary Table S5), necessitating further investigation to accurately assess their compaction phenotypes.

### Identification and validation of final hit strains

With 156 candidate strains identified, we initiated a stringent validation process to pinpoint the most promising hit strains (Fig. 3). The first step involved repeating the high-throughput experiment, with all candidate strains prepared and imaged in a common plate. Subjective evaluation of the resulting images (see Supplementary Material) excluded strains that exhibited DNA supercompaction, refining our list to 35 candidates for follow-up experiments.

**Figure 3.**
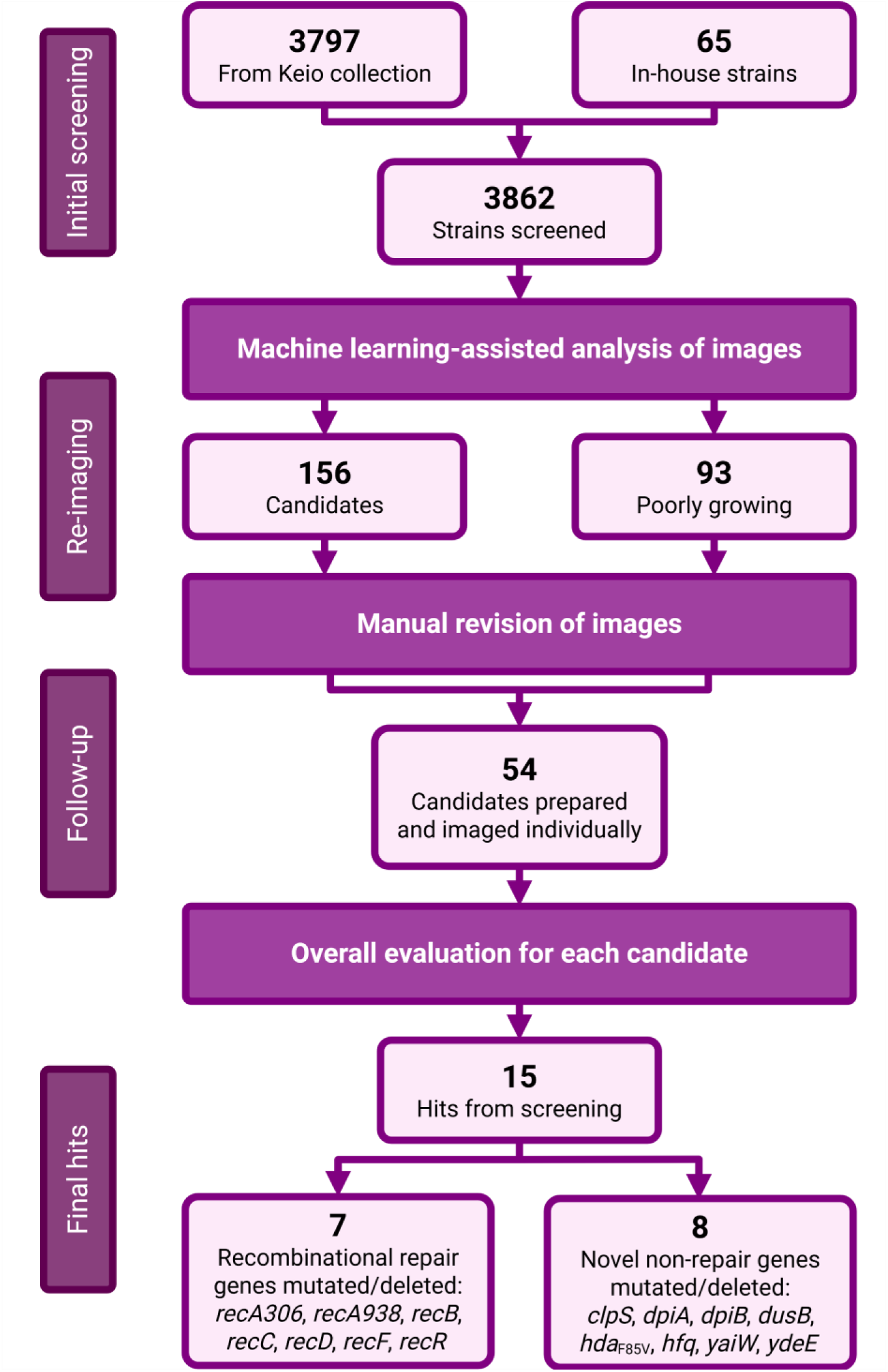
Flowchart depicting the selection process for identifying the hit strains from the screening. Following the initial screening of 3797 Keio collection strains and 65 in-house strains, 156 candidate strains were identified from the machine-learning assisted image analysis, along with 93 poorly growing strains. Both groups were reimaged using more controlled growth conditions, leading to 54 remaining candidates for follow-up validation experiments after subjective image evaluation. These experiments utilized both rich LB medium and supplemented AB minimal medium to individually culture strains prior to treatment and fixation. Manual imaging with a fluorescence microscope enhanced resolution and differentiation of compaction phenotypes. Following an overall evaluation, we ended up with the 15 hit strains listed. These comprised 7 strains with mutations or deletions of genes linked with recombinational repair and the SOS response, as well as 8 strains with mutations or deletions of novel genes not previously associated DNA compaction or repair. Created in BioRender (https://BioRender.com/wos5279).

For the 93 poorly growing strains, we performed a new replicate where all strains were cultured in individual tubes to ensure they reached exponential growth prior to CIP exposure, fixation, and DNA staining (see Supplementary Material). After high-throughput imaging, strains exhibiting DNA supercompaction were excluded, leaving 19 candidates for follow-up experiments. We were then left with a total of 54 candidate strains that appeared to have impaired DNA supercompaction responses (Supplementary Table S6).

Follow-up experiments were then conducted on the remaining 54 candidate strains to validate their compaction phenotypes and identify the final hits from the screening. To ensure precise evaluation, each strain was individually cultured to exponential growth and prepared for imaging on slides. This allowed us to use a fluorescence microscope with an oil immersion objective to improve resolution and better distinguish compaction phenotypes. Imaging was performed on samples grown in both rich LB medium—as in the screening— and in AB minimal medium. The minimal medium reduces the number of replicating chromosomes (37), which we expected would facilitate clearer distinction of compaction phenotypes. After thoroughly assessing these images alongside previously collected data for each candidate strain, those not clearly displaying distinct phenotypes from the wild-type control in the majority of cells examined were excluded, leaving us with a list of 15 final hit strains (Fig. 3). These hits comprised genes associated with recombinational repair, and novel genes not previously linked with DNA compaction or repair (details in Table 2). Four of the hit strains—*ΔdpiA*, *ΔdpiB*, *hda*_F85V_, and *ΔydeE*—demonstrated poor growth during experiments (Supplementary Table S5). Additionally, the *ΔdpiA* and *ΔdpiB* hit strains exhibited impaired supercompaction in only a subpopulation of filamenting cells, whereas the *ΔydeE* hit strain demonstrated notably short and small cells (Supplementary Fig. S2). With these final hits validated, we then turned to investigate the role of their mutations and deletions in the DNA supercompaction response.

**Table 2.** Details on the 15 hit strains with impaired DNA supercompaction from the screening, including compaction impairment levels categorized as major, moderate, or minor based on subjective evaluation. * Compaction impairment also includes situations where a proportion of cells have impaired compaction. ^a^ The SMG3 strain is a variant of the MG1655 wild type. ^b^ The *recA306* notation refers to the *Δ(srl-recA)306* mutation, which only leaves a small portion of the *recA* gene and can be considered a deletion of *recA* **(43–45)**.

| Strain | Genotype | Details about protein and gene | Compaction impairment* | Source |
| --- | --- | --- | --- | --- |
| JW0865 | BW25113<br>*ΔclpS* | Chaperone binding | Minor | Keio collection (23) |
| HI1733 | SC1148<br>*ΔdpiA* | SOS response induction after β-lactam exposure (59), DNA binding | Minor | In-house strain; (60) |
| HI1734 | SC1148<br>*ΔdpiB* | SOS response induction after β-lactam exposure (59) | Minor | In-house strain; (60) |
| JW3228 | BW25113<br>***ΔdusB*** | tRNA binding and modification, Co-transcribed with *fis* | Moderate | Keio collection |
| KS1115 | SMG3<sup>a</sup><br>***hda*<sub>F85V</sub>** | Inhibitor of re-initiation of DNA replication, Possible gain of function mutant (61) | Major | In-house strain |
| JW4130 | BW25113<br>***Δhfq*** | RNA-binding to stimulate RNA-RNA pairing | Moderate | Keio collection |
| STL9253 | MG1655<br>***recA306*<sup>b</sup>** | SOS response inducer after DNA damage, Homologous recombination protein, DNA binding | Major | In-house strain; (43–45) |
| ALS972 | MG1655<br>***recA938*** | SOS response inducer after DNA damage, Homologous recombination protein, DNA binding | Major | In-house strain; (62) |
| JW2788 | BW25113<br>***ΔrecB*** | Homologous recombination protein, DNA binding | Major | Keio collection |
| JW2790 | BW25113<br>***ΔrecC*** | Homologous recombination protein, DNA binding | Major | Keio collection |
| JW2787 | BW25113<br>***ΔrecD*** | Homologous recombination protein, DNA binding | Major | Keio collection |
| JW3677 | BW25113<br>***ΔrecF*** | Homologous recombination protein, DNA binding | Moderate | Keio collection |
| JW0461 | BW25113<br>***ΔrecR*** | Homologous recombination protein, DNA binding | Moderate | Keio collection |
| JW0369 | BW25113<br>***ΔyaiW*** | Unknown function | Minor | Keio collection |
| JW1527 | BW25113<br>***ΔydeE*** | Dipeptide exporter | Moderate | Keio collection |

### Hit strains exhibit delays in DNA supercompaction progression

In the first phase of our investigation, we explored how mutations and deletions in various hit strains affected DNA supercompaction progression. Our previous research established that RecN and RecA proteins are essential for proper supercompaction progression (18), prompting us to use the *ΔrecN* strain as a negative control for this investigation. To assess the progression rate of DNA supercompaction across the hit strains, we imaged samples fixed at 5-minute intervals, starting just before CIP exposure and continuing until 20 minutes post-exposure. DNA supercompaction progression was assessed through changes in DNA profile widths—the distribution of DNA along the cells’ long axis (Fig. 4A). We found that *recA306*, *recA938*, *ΔrecB*, and *ΔrecC* strains displayed impaired progression like the *ΔrecN* control, while *ΔrecD* and *ΔydeE* strains exhibited moderately impaired progression. The remaining hit strains showed minor delays in progression compared to the wild type. The DNA profile widths of these strains were distinguishable from the wild type at 10 and 15 minutes after exposure but became indistinguishable by 20 minutes (Fig. 4B).

**Figure 4.**
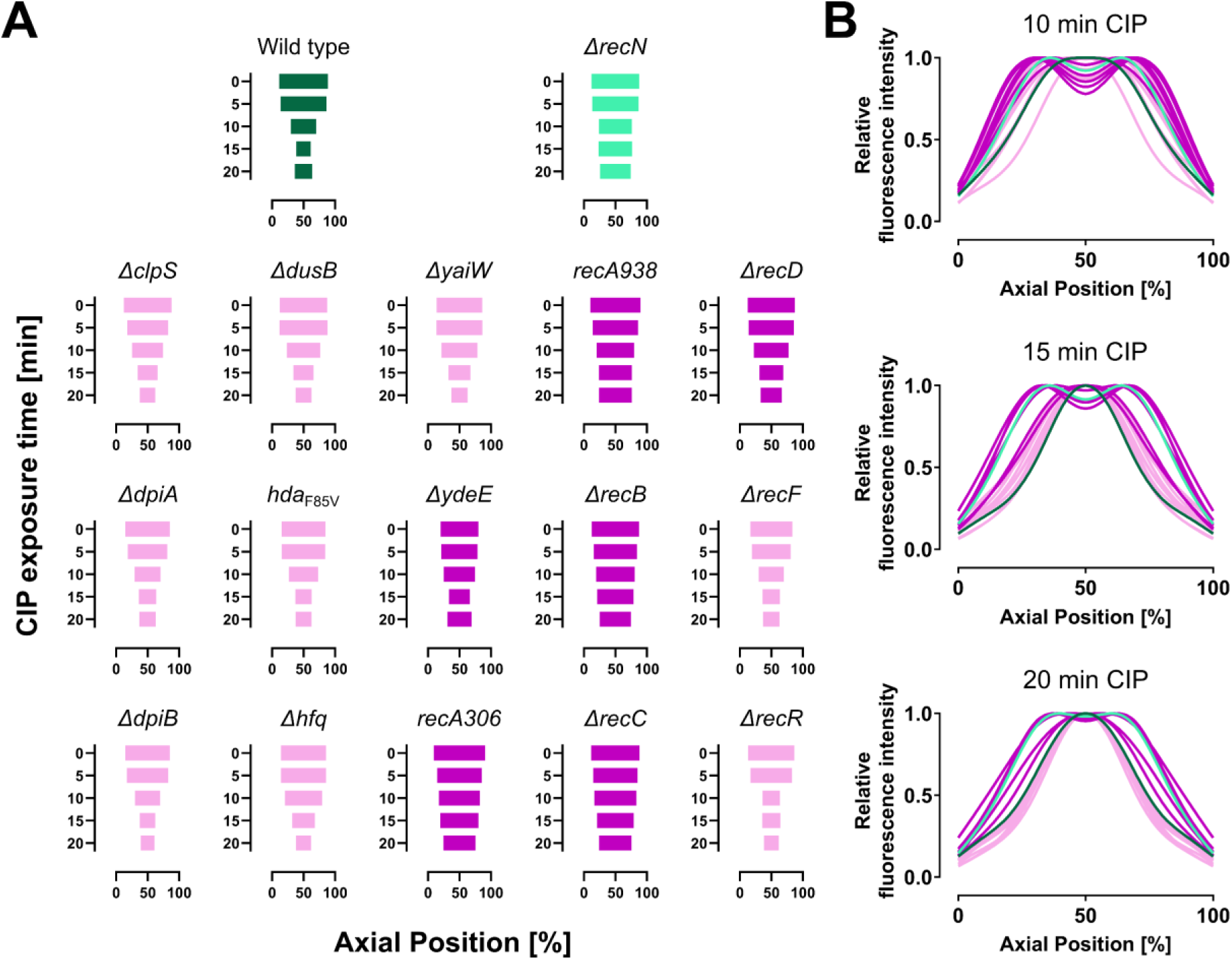
Compaction dynamics of hit strains and screening controls indicated by changes in DNA profile widths after various durations of CIP exposure. (**A**) Progression of compaction illustrated through profile widths for individual strains, captured at 5-minute intervals starting from just before CIP exposure to 20 minutes afterward. The DNA profile width is quantified relative to the cells’ long axis by measuring the distance between the outer bounds of symmetrical fluorescence peaks at 80% of the maximum averaged intensity at each time point. (**B**) Comparison of DNA profiles along the cells’ long axis for all strains after 10, 15, and 20 minutes post-CIP exposure. The wild-type control is presented in dark green, while *ΔrecN* is in light green. Dark magenta highlights strains with clear compaction impairment after 20 minutes of CIP exposure, and light magenta indicates strains with minor compaction impairment, mainly observable after 10 or 15 minutes of exposure. Data are from a single representative biological replicate.

### Genetic backgrounds influence how gene deletions affect supercompaction progression

As phenotypic effects of gene deletions often depend on genetic background through epistatic and compensatory effects (63–65), we examined whether a selection of gene deletions—*ΔdusB*, *Δhfq*, *ΔrecN*, and *ΔydeE*—originally screened in the BW25113 background, would also impair DNA supercompaction in other *E. coli* wild-type backgrounds. We transduced these deletions into the wild-type strains MG1655, AB1157, BL21 and CCUG17620 (Table 1). BW25113, the background of the Keio collection, is a 15-step descendant of the ancestral K-12 strain EMG2, while MG1655 is only two steps from EMG2 (23). The remaining strains represent a spectrum of *E. coli* genetic diversity, including the clinical isolate CCUG17620 (46–51).

As with hit strains, we cultured the various wild types and newly constructed gene deletion strains to exponential phase and fixed samples at 5-minute intervals from just before CIP exposure (10 µg/mL) until 20 minutes after. As earlier, we assessed DNA supercompaction progression through changes in DNA profile widths (Fig. 5A). All *E. coli* wild-type strains showed complete DNA supercompaction after 20 minutes of CIP exposure (Fig. 5A and B). Comparing compaction progression for deletion strains with the BW25113 strain revealed that each gene deletion clearly impaired progression in at least two genetic backgrounds, with considerable variations between backgrounds (Fig. 5A and B). The *ΔrecN* deletion consistently impaired supercompaction across all backgrounds, while the effects of *ΔdusB*, *Δhfq*, and *ΔydeE* deletions were more variable (Fig. 5A and B). In particular, the *Δhfq* deletion caused major delays in supercompaction progression in BL21 and CCUG17620 backgrounds, resembling the effects seen with *ΔrecN* deletions. Additionally, both *ΔdusB* and *ΔydeE* deletions moderately impaired progression in the CCUG17620 background (Fig. 5A and B).

**Figure 5.**
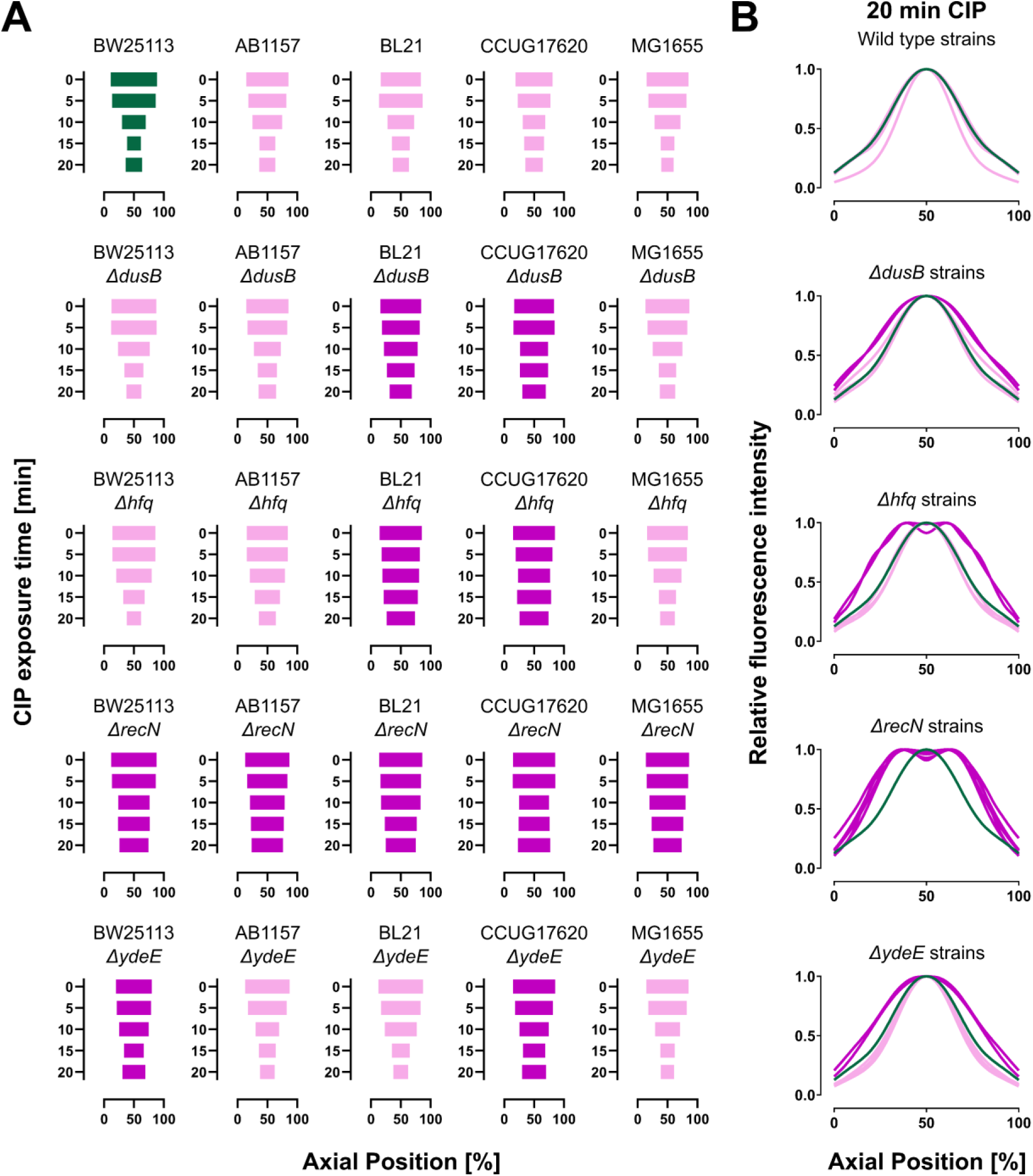
Influence of different *E. coli* wild-type backgrounds (BW25113, AB1157, BL21, CCUG17620, and MG1655) on compaction dynamics following deletions of *dusB*, *hfq*, *recN*, or *ydeE*. All strains were cultured to exponential phase in LB at 37°C. (**A**) Progression of compaction illustrated through profile widths for individual strains with various gene deletions across different backgrounds, captured at 5-minute intervals starting from just before CIP exposure to 20 minutes post-exposure. The DNA profile width is quantified relative to the cells’ long axis by measuring the distance between the outer bounds of symmetrical fluorescence peaks at 80% of the maximum averaged intensity at each time point. (**B**) Comparison of DNA profiles along the cells’ long axis for strains with identical gene deletions across different backgrounds after 20 minutes of CIP exposure. The screening wild-type strain (BW25113) is presented in dark green. Dark magenta highlights strains with clear compaction impairment after 20 minutes of CIP exposure, while light magenta indicates strains without observable compaction impairment at this exposure duration. Data are from representative biological replicates.

### Clinical isolates exhibit DNA supercompaction

A striking observation was that all transduced gene deletions resulted in impaired DNA supercompaction progression in the CCUG17620 background. CCUG17620, unlike typical lab-adapted *E. coli* strains, serves as a reference for clinical isolates (50). Given its clinical origins and susceptibility to DNA supercompaction impairment, we hypothesized that some clinical isolates might not initiate this response at all. To investigate whether DNA supercompaction is a common response to CIP exposure among various clinical isolates, we examined eight gram-negative clinical strains from *E. coli*, *Klebsiella oxytoca* and *Klebsiella pneumoniae,* isolated from blood, urine, bile fluid, and intestines. Cultured to exponential phase, samples from these strains were fixed just before and 20 minutes after CIP exposure (10 µg/mL). Analyzing the DNA distribution within the cells revealed that all clinical isolates clearly exhibited DNA supercompaction following 20 minutes of CIP exposure (Fig. 6).

**Figure 6.**
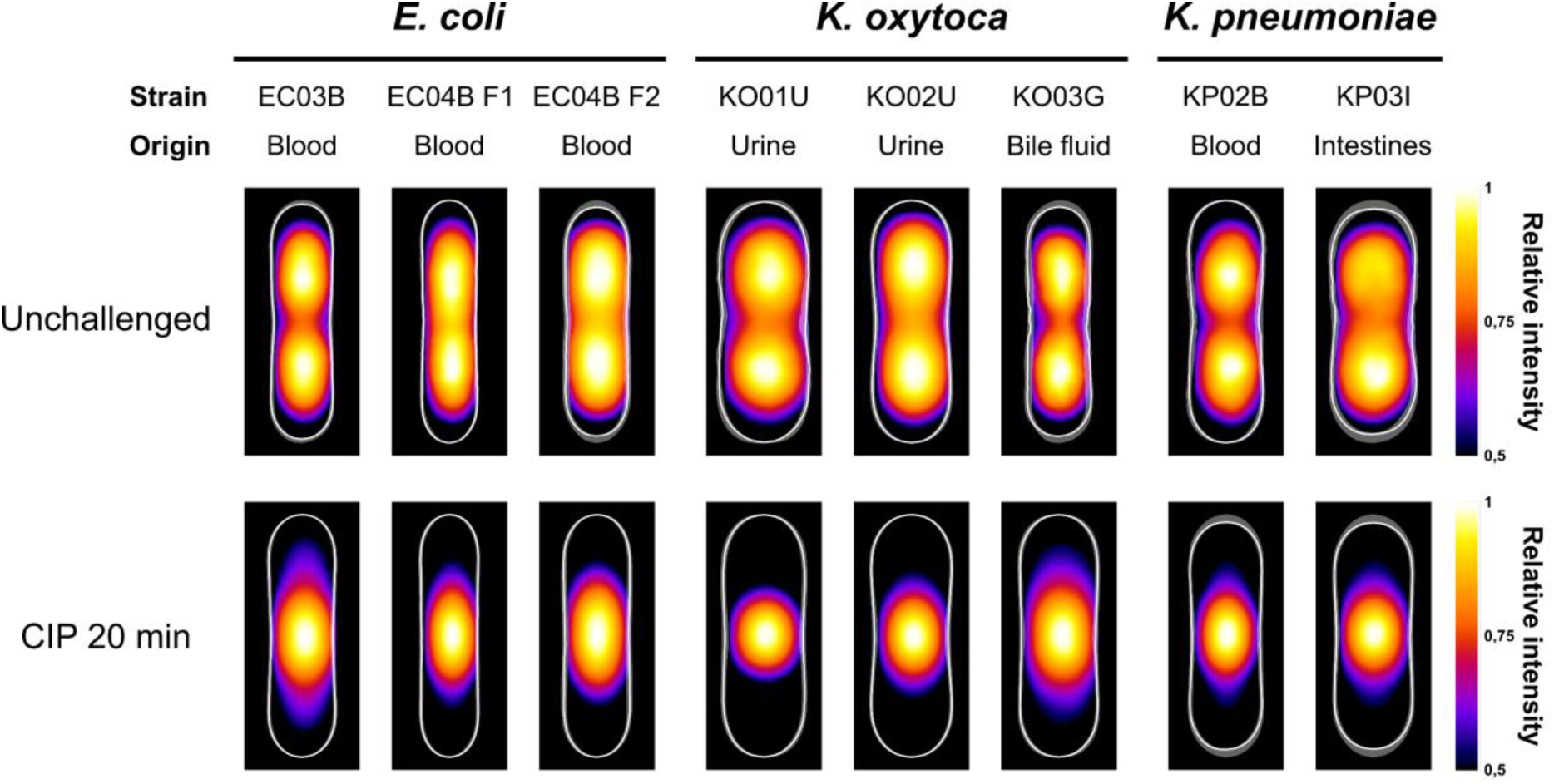
DNA supercompaction occurs in clinical strains following CIP exposure. The strains include *E. coli*, *K. oxytoca*, and *K. pneumoniae*, isolated from blood, urine, bile fluid, and intestines. Kymograph heat maps display the Hoechst 33258 relative intensity distribution in unchallenged cells (top row) and cells exposed to 10 µg/mL CIP for 20 minutes (bottom row). Results are from representative biological replicates and are averaged from 37 to 1395 cells per sample. EC04B F1 and EC04B F2 are distinct strains from the same patient.

### SOS response activity varies between hit strains

The DSBs caused by CIP exposure trigger the SOS response (6), which regulates expression of RecN and RecA—proteins essential to proper progression of DNA supercompaction (18). RecN has a particularly large increase in expression at the initiation of the SOS response (8, 9), coinciding with the timing of the supercompaction response (18). To assess whether the compaction phenotypes of hit strains resulted from a reduced SOS response, and thus decreased levels of RecN or RecA, we quantified their SOS response activities. All strains were transformed with the p*lexA*-*gfp* reporter plasmid, enabling *lexA* promoter-regulated GFP expression measurable by flow cytometry (40, 41). Strains cultured to exponential phase were sampled just before and at 30-minute intervals after CIP exposure (10 µg/mL) to measure GFP fluorescence in 20,000 cells per sample. Sampling continued until 120 minutes post-exposure, when fluorescence signals plateaued in the wild type and other strains (Supplementary Fig. S3A and B). To account for filamentation and presumably higher GFP levels in larger cells, we normalized mean GFP fluorescence signals to the mean forward scatter for each sample.

As expected due their impaired processing of DSBs (6, 66), *recA*, *recB*, and *recC* deletion strains failed to increase their SOS response activities immediately after CIP-induced DNA damage (Fig. 7A) and maintained very low activity levels 30 minutes post-exposure (Supplementary Fig. S3C). In contrast, the *ΔrecD* strain showed only a minor increase in SOS response activity (Fig. 7A) yet achieved activity levels similar to wild type at 30 minutes post-exposure (Supplementary Fig. S3C), consistent with its ability to load RecA despite compromised regulation (66). Among novel hit strains, while some exhibited slightly impaired SOS response increases at this early stage, none of these differed significantly from the wild type (Fig. 7B). Notably, the *ΔdusB* and particularly the *ΔyaiW* strains displayed pronounced increases in SOS response activity levels during the first 30 minutes of CIP exposure (Fig. 7B) and achieved significantly higher levels than the wild type at this stage (Supplementary Fig. S3D). At 120 minutes post-exposure, *recA*, *recB*, and *recC* deletion strains remained at very low activity levels, whereas the *ΔrecD* strain sustained levels similar to the wild type (Fig. 7C). The *ΔdusB* and *ΔyaiW* strains continued to display very high SOS response activities, indicating sustained elevations of DNA damage responses (Fig. 7D). Most other strains reached activity levels similar to the wild type, except for the *ΔrecN* strain, which showed a small yet significant reduction in activity at 120 minutes post-exposure (Fig. 7C).

**Figure 7.**
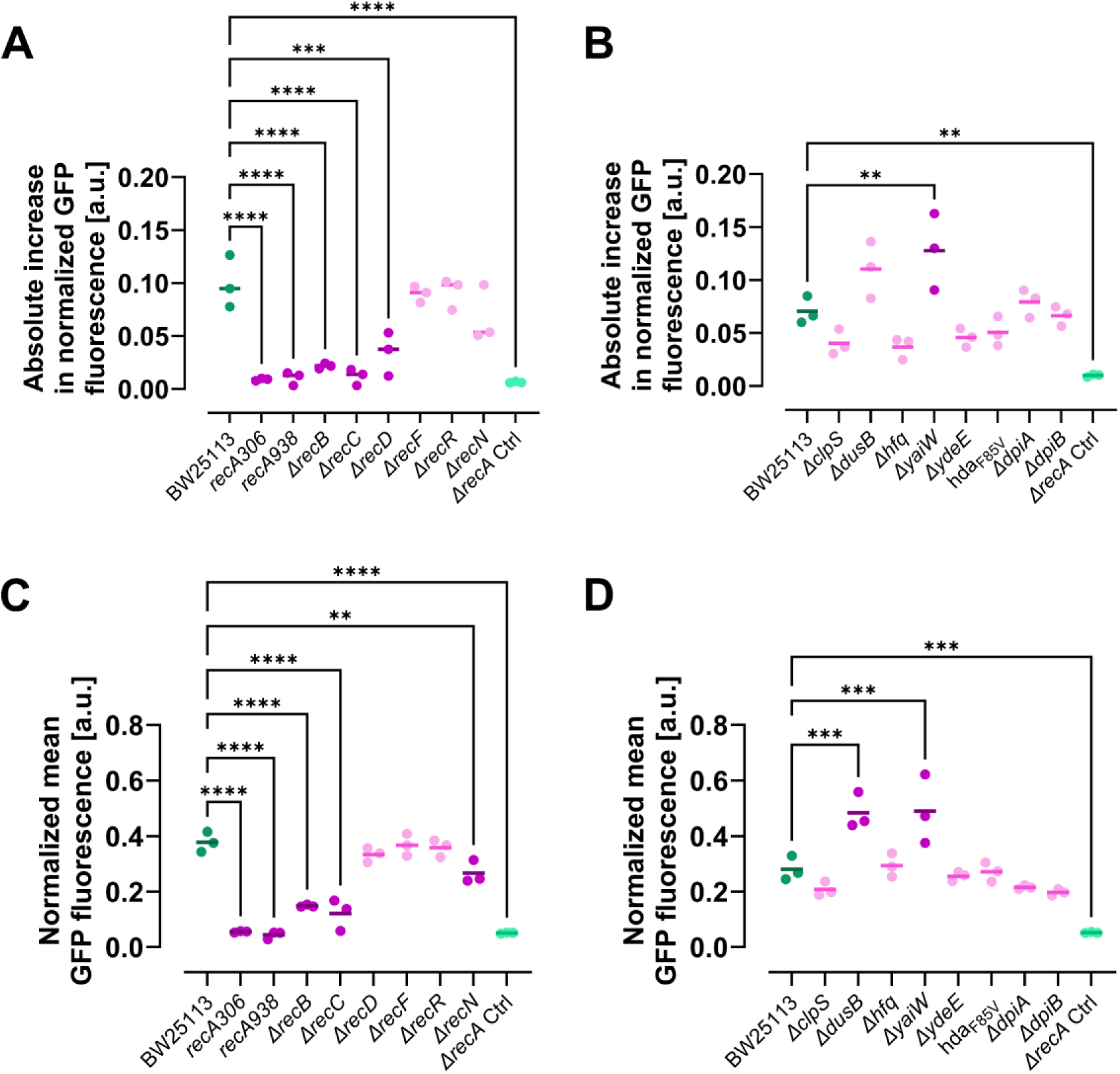
Quantification of SOS response activity in hit strains after exposure to 10 µg/mL CIP. GFP fluorescence, regulated by the *lexA* promoter and expressed from a reporter plasmid, indicates SOS response activity and was measured in 20,000 cells per sample using a flow cytometer. (**A** and **B**) show the change in SOS response activity from baseline (0 min) to 30 minutes post-CIP exposure, while (**C** and **D**) present activity measured 120 minutes after exposure. The assay was performed separately for strains with mutations or deletions of (**A** and **C**) genes linked to recombinational repair and the SOS response, and (**B** and **D**) genes not previously associated with DNA compaction or repair. A *ΔrecA* strain from the Keio collection (JW2669) served as negative control. SOS response activity was quantified by normalizing mean GFP fluorescence to mean forward scatter area (mean cell size). Lines represent means from three biological replicates; dots indicate individual replicate means. The wild type (BW25113) is presented in dark green, *ΔrecA* in light green. Strains with significantly different activity from the wild type of the corresponding set of experiments are in dark magenta; others are in light magenta. Ordinary one-way ANOVA with Dunnett correction was used for comparisons; only significant differences are annotated. \*\**P* ≤ 0.01; \*\*\**P* ≤ 0.001; \*\*\*\**P* ≤ 0.0001.

### RecN distribution and nucleoid colocalization is affected by novel gene deletions in select hit strains

Next, we explored whether the impaired supercompaction observed in the hit strains with novel gene deletions could be attributed to altered RecN dynamics. For this investigation, we compared the nucleoid compaction and RecN dynamics of *ΔclpS*, *ΔdusB*, *Δhfq*, and *ΔyaiW* strains against the wild type using live-cell imaging. We chose not to include the remaining four hit strains due to distinct characteristics that could affect analyses. Specifically, we omitted the *hda*_F85V_ strain owing to its very slow growth, *ΔydeE* due to its small cell size, and *ΔdpiA* and *ΔdpiB* since impaired supercompaction occurred only in filamenting subpopulations (Supplementary Fig. S2).

To visualize the cells’ DNA during live-cell imaging, we transduced an HU-mCherry construct into the strains with BW25113 backgrounds (18, 52). Additionally, we introduced the pSOS plasmid to enable GFP-RecN expression at native RecN levels (17, 18). Cells were cultured to exponential phase, exposed to 10 µg/mL CIP, and after a 1-minute incubation, prepared for imaging on agar pads. Imaging began 10 minutes post-CIP exposure using live-cell spinning disk microscopy.

In our prior publication (18), we demonstrated that wild-type cells (Fig. 8A) exhibit rapid DNA supercompaction, with GFP-RecN foci clustering near the compact nucleoid. In the current investigation using live-cell imaging, the *ΔyaiW* strain showed substantial impairment in its supercompaction response (Fig. 8B, upper panel). In contrast, *ΔdusB* and *Δhfq* strains displayed only minor delays in supercompaction progression, while the *ΔclpS* strain exhibited no observable delays (Fig. 8B, upper panels). Strikingly, GFP-RecN had more dispersed localization in these hit strains compared to the wild type (Fig. 8B vs A, lower panels). Despite the increased dispersion, the number of GFP-RecN foci for the hit strains remained at similar levels to the wild type (Fig. 8C). The dispersed GFP-RecN distribution in hit strains was also reflected in a lower colocalization with the nucleoid compared to the wild type (Fig. 8D), which was particularly pronounced in the *ΔyaiW* strain.

**Figure 8.**
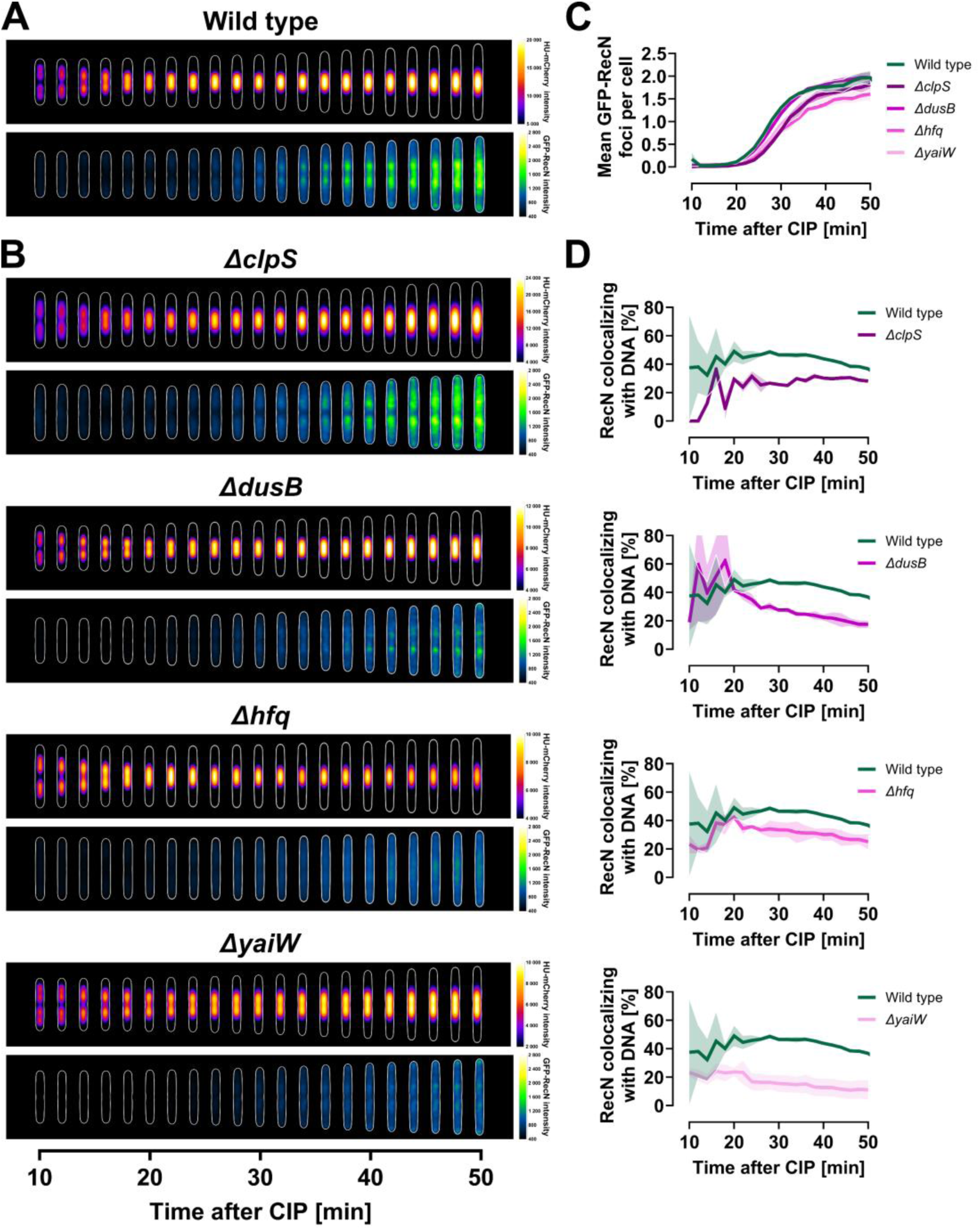
Live-cell analysis of compaction and GFP-RecN dynamics after CIP exposure for wild type and select hit strains—*ΔclpS*, *ΔdusB*, *Δhfq*, and *ΔyaiW*. All strains contained the pSOS plasmid for GFP-RecN expression, were cultured in LB at 37°C, and imaged at 2-minute intervals using live-cell spinning disk microscopy starting 10 minutes after CIP exposure. Data for the wild-type strain (A, C, and D) were previously published in **(3)**. (**A** and **B**) Kymograph heat maps displaying intensity distribution for HU-mCherry (upper panels) and GFP-RecN (lower panels) within cells following CIP exposure. Note the variations in HU-mCherry intensity scales. Results are from representative biological replicates and are averaged from 328-480 cells for wild type, 105-190 cells for *ΔclpS*, 235-381 cells for *ΔdusB*, 958-1807 cells for *Δhfq*, and 357-726 cells for *ΔyaiW*. (**C**) Mean number of GFP-RecN foci per cell versus time post-CIP exposure for all strains. (**D**) Percentage of GFP-RecN foci colocalizing with DNA over time post-CIP exposure, comparing individual hit strains to wild type. Lines represent means from two biological replicates, with shaded regions indicating SEM. Data in the early time points of (D) only reflect a small number of cells with very weak foci presence, reflected by the large SEM.

### Only some hit strains exhibit increased sensitivity to ciprofloxacin

Our research previously demonstrated that a *ΔrecN* strain is extremely sensitive to CIP, with survival dropping to undetectable levels after only 1 minute of 10 µg/mL CIP exposure (18). Given that *ΔrecA* strains are also known for their heightened sensitivity to DSBs (28, 67–69), we explored whether similar sensitivity extends to other hit strains. We tested the survival of selected hit strains after various durations of exposure to 10 µg/mL CIP. The *hda*_F85V_ strain was not included due to very poor growth, while the *ΔdpiA* and *ΔdpiB* strains were omitted because of the subpopulation characteristics observed when assessing their DNA supercompaction response (Supplementary Fig. S2).

Similar to *ΔrecN*, *ΔrecB* and *ΔrecC* strains exhibited extreme sensitivity to CIP (Fig. 9A, *P*-values <0.0001, Supplementary Table S7), reflecting their inability to initiate DNA repair by processing DSBs (66). While *ΔrecR* resembled the wild type (*P*=0.3252), *ΔrecD* and *ΔrecF* strains displayed reduced sensitivity compared to the wild type (Fig. 9A, *P*-values <0.0001, Supplementary Table S7). Among novel hits, *ΔdusB* exhibited the highest sensitivity (Fig. 9B). The *ΔclpS* and *ΔyaiW* strains appeared slightly more sensitive than wild type, while *Δhfq* and *ΔydeE* appeared slightly less sensitive, though none of these differences from wild type among novel hit strains were statistically significant (*P*-values 0.07-0.16, Supplementary Table S8).

**Figure 9.**
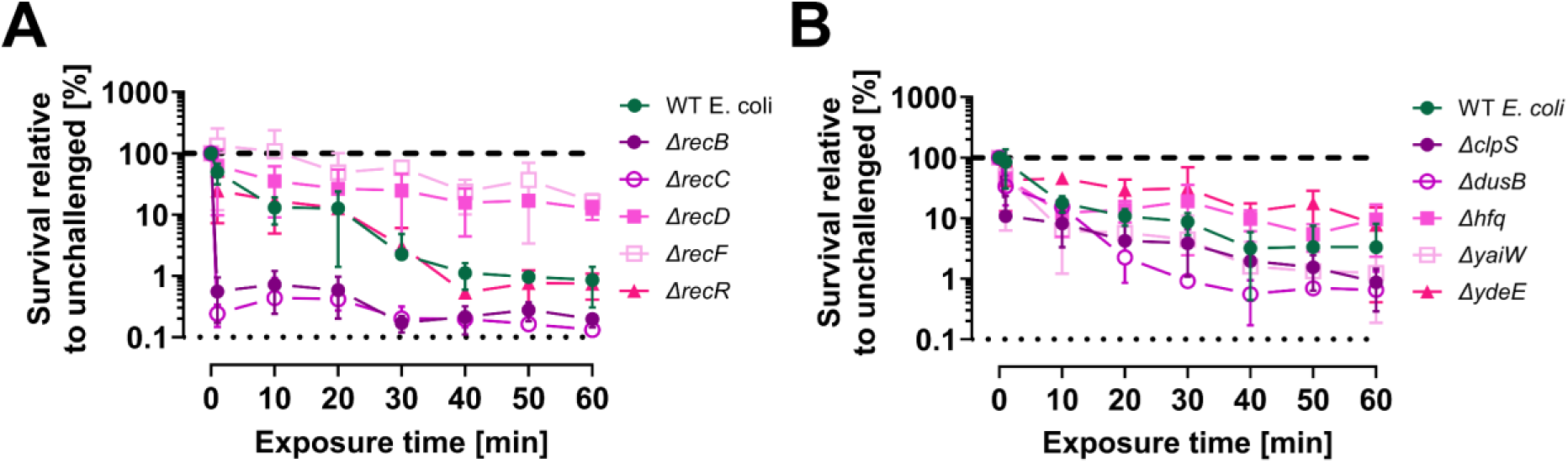
CIP sensitivity varies across different hit strains. We assessed time-dependent survival after exposure to 10 µg/mL CIP for hit strains with gene deletions (**A**) already linked to recombinational repair and the SOS response, and (**B**) not previously associated with DNA compaction or repair. Samples were collected just before CIP exposure and at time points ranging from 1 to 60 minutes afterwards. Exposure time refers to the duration of CIP exposure before washing and plating the cells. Survival was measured as colony forming units (CFU) per mL, with relative survival calculated through comparison with unchallenged parallels. Thick dashed lines indicate survival of unchallenged parallels; thin dotted lines mark the assay’s detection limit. Symbols with connecting lines represent means from 3-4 biological replicates, with error bars indicating standard deviations.

### UV-induced DNA compaction and survival: major impairment observed only with *recA* deletions

RecN has been found to influence DNA compaction following both CIP and UV exposure (18, 19). While both treatments are genotoxic, CIP primarily induces DSBs (2, 3), whereas UV irradiation generates bulky adducts resulting in single-strand gaps in DNA (70, 71). UV-induced DNA compaction has been observed with irradiation levels as low as 3 J/m^2^ in *E. coli* (19). Here, we studied the effect of 5 J/m^2^ UV irradiation on compaction and survival in exponentially growing strains (Fig. 10).

**Figure 10.**
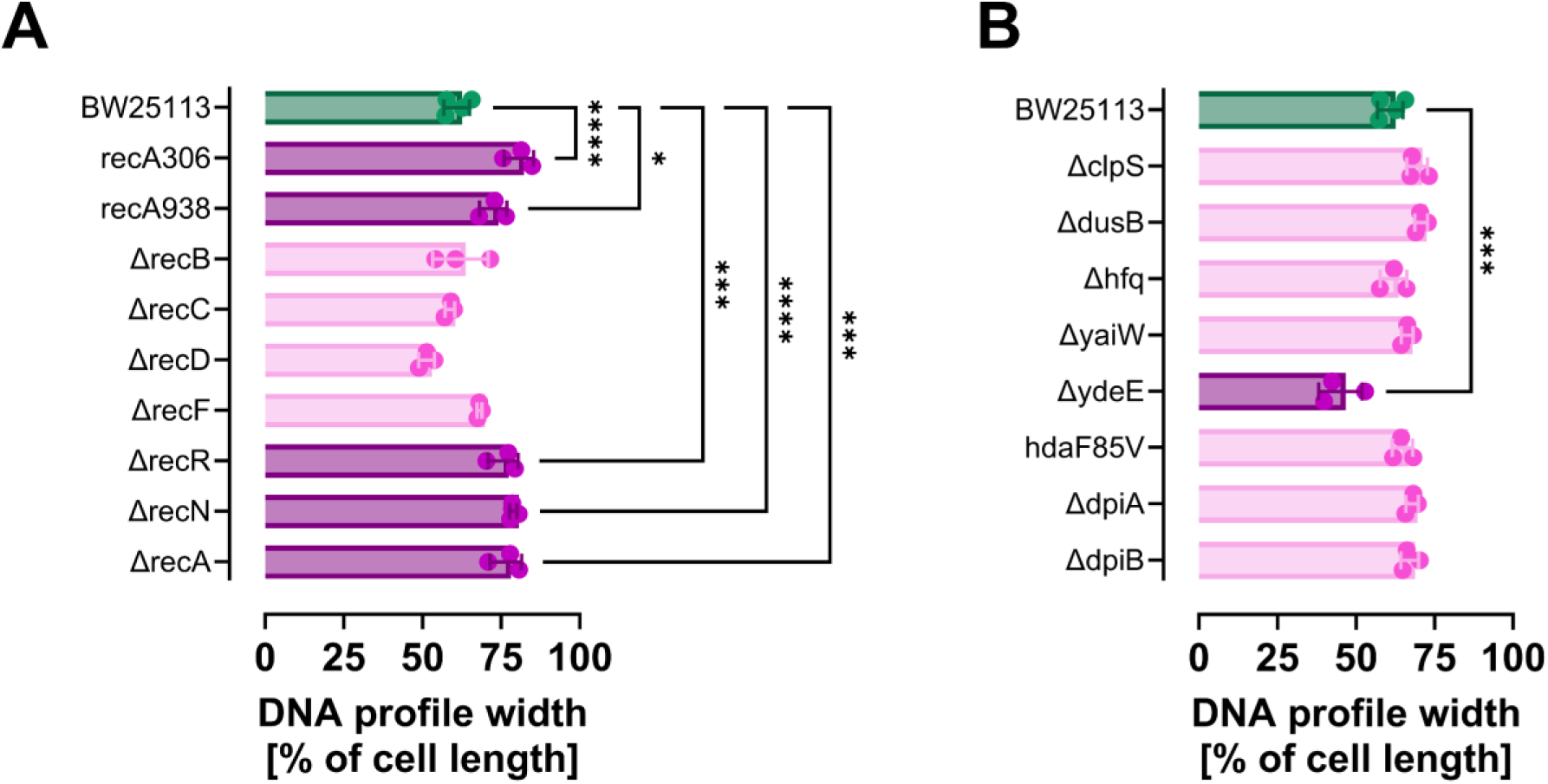
Some hit strains exhibit impaired DNA compaction following UV exposure. The effect of mutations and deletions on DNA compaction was assessed by comparing the DNA profile widths of hit strains to the wild type (BW25113) after 5 J/m^2^ UV irradiation and 15-minute recovery. Profile widths were quantified relative to the cells’ long axis by measuring the distance between the outer bounds of symmetrical fluorescence peaks at 80% of the maximum averaged intensity. Results are presented for hit strains with mutations or deletions of (**A**) genes linked to recombinational repair and the SOS response, including a *ΔrecN* strain (JW5416), and (**B**) genes not previously associated with DNA compaction and repair. Columns represent means from 3-4 biological replicates; dots show individual replicate values; error bars indicate standard deviation. The wild type is shown in dark green. Strains with significantly different DNA profile widths from the wild type appear in dark magenta, while others are in light magenta. Comparisons used ordinary one-way ANOVA with Dunnett correction; only significant differences are annotated. \**P* ≤ 0.05; \*\*\**P* ≤ 0.001; \*\*\*\**P* ≤ 0.0001.

To assess UV-induced DNA compaction, strains were UV irradiated, incubated 15 minutes for recovery, followed by fixation and staining with Hoechst 33258. We then imaged the stained samples using fluorescence microscopy and quantified DNA profile widths to evaluate the progression of compaction (Fig. 10), consistent with earlier methods. In accordance with prior research (19), strains with *recA*, *recN* or *recR* deletions failed to exhibit DNA compaction following UV irradiation, unlike the wild type (Fig. 10A). Although the *ΔrecF* strain showed a small delay in compaction progression, this delay was unexpectedly not statistically significant compared to the wild type (Fig. 10A), despite RecF’s role in the RecFOR pathway responsible for initiating repair of single-strand gaps (72). Among most novel hit strains, there was an observable yet non-significant tendency of delayed compaction progression (Fig. 10B). Surprisingly, the *ΔydeE* strain demonstrated clearly enhanced DNA compaction post-UV irradiation (Fig. 10B), though this result was likely an artefact due to the strains’ small cell size (Supplementary Fig. S2).

Sensitivity to UV irradiation was assessed by comparing the relative survival of irradiated samples to unchallenged parallels for each strain. The wild type exhibited a survival rate averaging 74% at the 5 J/m^2^ dose (Fig. 11). Among all tested strains, only those with *recA* deletions showed significantly higher sensitivity to this UV dose compared to the wild type (Fig. 11A and B). Despite its impaired UV-induced compaction, the *ΔrecN* strain did not exhibit any difference in survival from the wild type at the 5 J/m^2^ dose (Fig. 11A). To investigate whether differences in UV sensitivity might emerge at higher doses—where compaction or the presence of RecN could be more critical for survival (18)—we compared dose-dependent survival rates between the *ΔrecN* and wild-type strains across doses ranging from 10 to 70 J/m^2^. In contrast to its heightened sensitivity to CIP and other DSB-inducing agents (14, 15, 17, 18, 28), the *ΔrecN* strain showed only a small, non-significant increase in UV sensitivity compared to wild type across the tested dose range (Fig. 11C, *P*=0.0603, Supplementary Table S9). These results align with earlier research (14, 73, 74) and imply that RecN is primarily involved in DSB repair rather than in single-strand gap repair.

**Figure 11.**
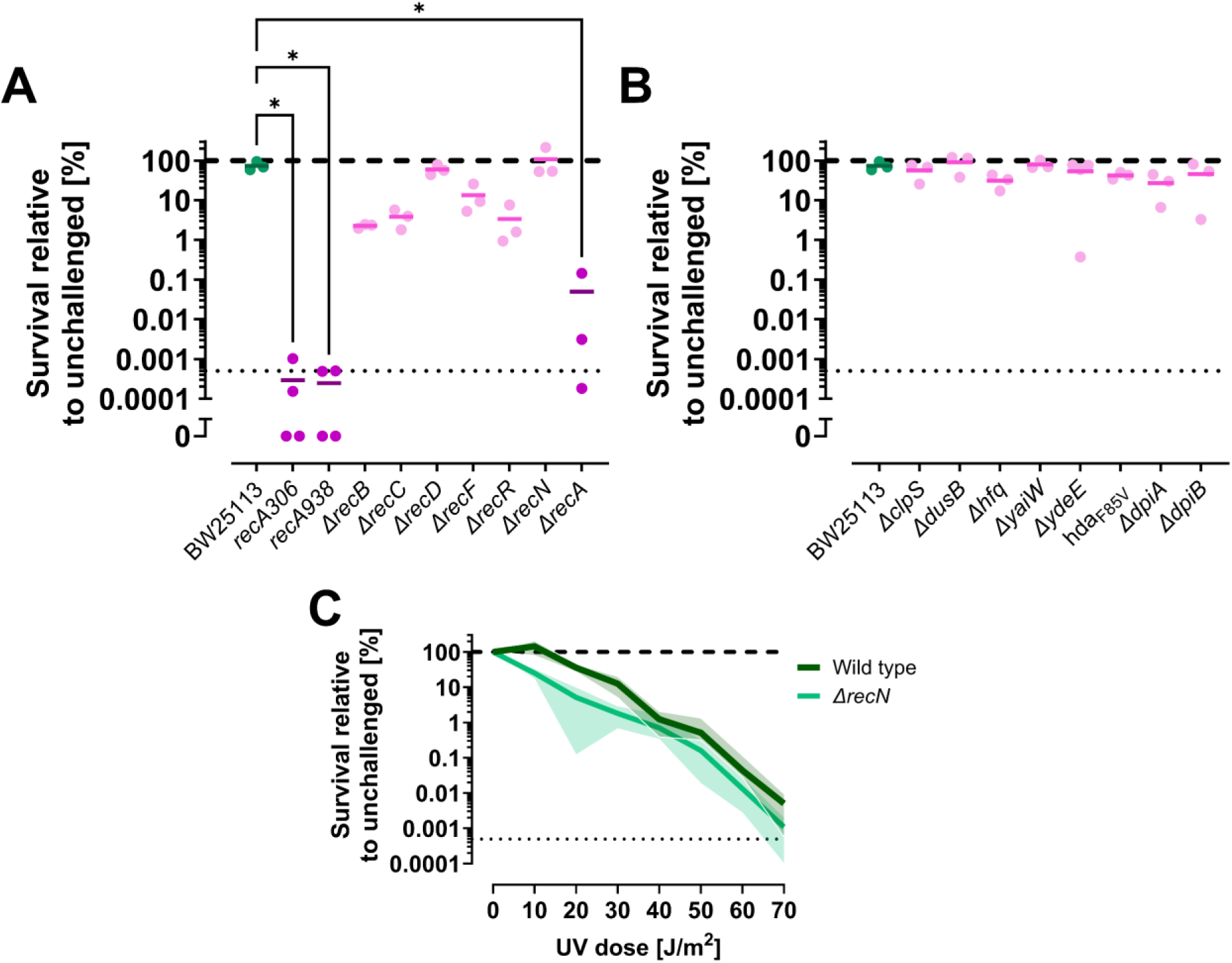
UV sensitivity varies among different hit strains, while the *ΔrecN* strain (JW5416) displays sensitivity similar to the wild type (BW25113). (**A** and **B**) Survival after 5 J/m^2^ UV irradiation was assessed for hit strains with mutations or deletions of (**A**) genes already linked to recombinational repair and the SOS response, including the *ΔrecN* strain, and (**B**) genes not previously associated with DNA compaction or repair. (**C**) Dose-dependent survival of *ΔrecN* versus wild type evaluated across UV doses ranging from 10 to 70 J/m^2^. Survival was measured as colony forming units (CFU) per mL, with relative survival calculated through comparison with unchallenged parallels. (A and B) Solid lines indicate means from 3-4 biological replicates; dots represent means from individual replicates. The wild type is shown in dark green. Strains with significantly different UV sensitivity from the wild type appear in dark magenta; others are in light magenta. Comparisons were made using ordinary one-way ANOVA with Dunnett correction; only significant differences are annotated. (C) Solid lines represent means from 3-4 biological replicates; shaded regions indicate standard deviation. Thick dashed lines show survival of unchallenged parallels; thin dotted lines mark the assay’s detection limit. \**P* ≤ 0.05.

## DISCUSSION

In the current investigation, we aimed to identify genetic factors involved in DNA supercompaction, prompted by our recent discovery that RecN and RecA are essential for this process in *E. coli* following severe DNA damage (18). To achieve this goal, we developed a strategy comprising high-content imaging, machine learning-assisted classification of compaction phenotypes, and a stringent validation process (Fig. 1). This strategy enabled us to effectively screen the CIP-induced compaction phenotypes for the entire Keio collection of *E. coli* single-gene deletion strains alongside additional in-house strains. Consequently, we identified 15 hit strains featuring deletions of both known and novel genetic factors (Table 2) that influence CIP-induced DNA supercompaction in *E. coli* (Fig. 4). The effects of these deletions varied across different genetic backgrounds (Fig. 5). Furthermore, DNA supercompaction was observed after CIP exposure in various gram-negative clinical isolates (Fig. 6). Notably, the supercompaction process was impaired for all hit deletions transduced into the clinical reference background CCUG17620 (Fig. 5), indicating these genetic factors may be especially relevant for DNA supercompaction in clinical settings. Although most novel gene deletions did not significantly alter SOS response activity (Fig. 7), select hit deletions caused GFP-RecN to become more dispersed within the cell and display reduced colocalization with the nucleoid after DNA damage (Fig. 8), potentially contributing to impaired DNA supercompaction dynamics. Despite impaired compaction following both CIP exposure (Fig. 4) and UV irradiation (Fig. 10), only a subset of hit strains exhibited heightened sensitivity to these genotoxic agents (Fig. 9 and 11, respectively).

### Screening for imageable phenotypic differences: limitations of the high-throughput approach

We utilized the Keio collection of single-gene deletion strains to uncover genetic factors influencing DNA supercompaction. While this collection is a highly valuable resource when screening for phenotypic differences caused by single-gene deletions, it also comes with several inherent limitations (24, 25, 63). First, since essential genes cannot be deleted due to lethality (23), the collection includes only strains with non-essential gene deletions, thereby limiting the discovery of essential genes involved in DNA supercompaction. Second, deleting specific non-essential genes may impose strong selective pressure on strains, potentially leading to confounding mutations. These mutations can cause unrelated phenotypic differences and complicate interpretation of gene deletion effects (63). Furthermore, growth conditions might drive the acquisition of secondary mutations that further confound phenotypic conclusions (75). Third, functional redundancy among genes can mask phenotypes in single-gene deletion strains and conceal the involvement of specific genes in DNA supercompaction. Fourth, inherent variations in growth rates among strains complicate the synchronization of growth phases and cellular densities, both of which are critical requirements for effective genome-wide screening (28, 30, 58). Finally, because the Keio collection is restricted to the genetic background of the *E. coli* BW25113 strain—a lab-adapted, 15-step descendant of the ancestral K-12 strain EMG2 (23)—epistatic and compensatory effects may arise, which could mask the effect of certain gene deletions on DNA compaction and limit the applicability of our findings to diverse bacterial populations.

To process the nearly 50,000 images captured during our investigations, we relied on machine learning-assisted image analysis. While this approach provided objective and quantitative analysis (35), some limitations warrant consideration when interpreting our results. First, the machine learning model’s performance relies heavily on training data quality (35, 76–78). Preparing the training dataset required manual annotation, which is both time-consuming and prone to human error. Although we strived to create a diverse and representative dataset, any biases present in these data would inevitably propagate to the output classifications (35, 76–78). Second, phenotype classification accuracy depends on image quality. Our high-throughput approach, while efficient, used automatic focusing set across the plate bottom, which may have been suboptimal for some wells. Even minimal blurring from suboptimal focus could hinder distinction of nucleoid lobes, making it difficult to accurately classify compaction phenotypes.

Our Phenotype model effectively identified strains with normal DNA supercompaction, although its performance was suboptimal for those exhibiting *ΔrecN* and unchallenged phenotypes (Fig. 2B). Confusion between phenotypes appeared to primarily result in unclassified samples, though it also led to the unavoidable false predictions expected from classification models, potentially leading us to miss some hits in the screening. In an attempt to account for the confusion, we first evaluated the replicates matching the selection criteria (Supplementary Material), and then ensured all replicates were evaluated for strains with at least one matching replicate. Our stringent validation process aimed to filter out strains falsely predicted to have impaired supercompaction or confounded by the high-throughput growth conditions. Despite our efforts, the *ΔrecA* strain from the Keio collection was notably absent from the candidate list, presumably due to the high-throughput growth conditions and suboptimal focus for its images. Subsequent investigations showed that *recA* deletion strains, including the Keio collection strain, consistently failed to compact their nucleoids after 20 minutes of CIP exposure (Supplementary Fig. S4). Though the screening may not have uncovered all genetic factors, it successfully identified multiple genes that influence DNA supercompaction (Table 2), paving the way for a better understanding of this process.

### Novel hit genes and their potential functions in DNA supercompaction

Several genes identified from our screening have not previously been linked to DNA compaction or repair, prompting us to explore their potential involvement in DNA supercompaction using insights from our experiments and other existing knowledge.

Among these, the *hfq* gene deletion substantially impaired DNA supercompaction in BL21 and CCUG17620 backgrounds (Fig. 5), although it only caused minor delays in the BW25113 background (Fig. 4). Some of these effects may be ascribed to a slightly weakened SOS response induction (Fig. 7B) associated with reduced RecN expression (Fig. 8B). Hfq, beyond its primary role as a chaperone for sRNA-mRNA interactions, has been found to interact with DNA, albeit with lower affinity than RNA (79). Interestingly, deletion of *hfq* has recently been implicated in reducing CIP uptake (80)—an effect potentially linked to Hfq’s influence on genes involved in regulation of pH homeostasis, which may affect the pH-dependent CIP flux through porins or across the lipid bilayer (80–82). Despite a potential reduction in CIP uptake, we did not observe any substantial effect on survival (Fig. 9B). Such findings indicate that Hfq might influence DNA supercompaction through mechanisms other than CIP uptake, warranting further research into the exact mechanisms and how Hfq affects SOS response induction.

Among the remaining novel hit genes, the *ydeE* deletion caused the clearest delays in supercompaction progression within the BW25113 background (Fig. 4). Both *ydeE* and *dusB* deletions impaired supercompaction in the CCUG17620 background, while their influence on the response varied in other genetic backgrounds (Fig. 5). While the *ΔydeE* strain had similar SOS response activity to wild type, the *ΔdusB* and *ΔyaiW* hit strains showed substantial increases in SOS activity relative to wild type (Fig. 7B and D). Surprisingly, this heightened SOS activity was not reflected in the expression of RecN (Fig. 8B vs A) but was associated with a notable drop in RecN colocalization with DNA (Fig. 8D). Furthermore, *ΔyaiW* exhibited considerable supercompaction impairment in live-cell imaging compared to other novel hit strains (Fig. 8B). The DusB protein is recognized as a tRNA-modifying enzyme influencing translation (83, 84), whereas *yaiW* and *ydeE* genes are predicted to encode outer (85) and inner (86) membrane-associated proteins, respectively. Both *yaiW* and *ydeE* are regulated by CpxR (86, 87), while *yaiW* is also part of the σ^E^ regulon (88). CpxR and σ^E^ are key regulators of membrane stress responses, indicating the involvement of YaiW and YdeE in these cellular responses. Moreover, Hfq proteins have been shown to primarily localize to the inner membrane (89), suggesting a potential interaction with membrane-associated proteins. Additionally, *dpiA* and *dpiB* genes are involved in SOS response induction through a two-component signal following exposure to β-lactams that challenge membrane integrity (59); subpopulations of filamenting cells lacking these genes displayed impaired supercompaction (Supplementary Fig. S2). Our group has also previously revealed that a strain lacking DinQ, an inner membrane protein, exhibits slightly impaired DNA compaction following UV irradiation (90). Collectively, these findings point toward a complex interplay between DNA compaction dynamics and membrane-associated proteins, underscoring the need for deeper exploration of the underlying molecular mechanisms.

### Updated understanding of DNA supercompaction as a clinically relevant response

Our current investigation has reinforced our understanding of RecN and RecA proteins as the key orchestrators of DNA supercompaction (18), despite operating within an environment of potentially undiscovered mechanisms. Strains lacking these proteins showed impaired supercompaction progression (Fig. 4), aligning with prior findings (18), and *recN* deletion consistently hindered supercompaction across all tested genetic backgrounds (Fig. 5). Additionally, strains lacking RecB, RecC, and RecD exhibited impaired DNA supercompaction responses (Fig. 4). The RecBCD complex is important for processing DSBs and loading RecA for SOS response initiation and subsequent homologous recombination (6, 66). Deleting any of these four proteins diminished SOS response initiation (Fig. 7A) and resulted in consistently low SOS activity in all strains except the *ΔrecD* strain (Fig. 7C and Supplementary Fig. S3A), which is known to exhibit RecA loading but in a dysregulated manner (66). Furthermore, findings from novel hit strains imply that SOS response activity is not predictive of supercompaction progression (Fig. 4 and 7D), in accordance with our previous findings that RecA contributes to DNA supercompaction beyond SOS response induction (18). Similarly, the RecBCD proteins may be important for DNA supercompaction by facilitating RecA’s DNA binding and efficiency in the process beyond SOS induction, or through mechanisms yet to be discovered.

While DSBs rely on RecBCD processing, UV-induced single-strand gaps require the RecFOR proteins for RecA loading and subsequent SOS response initiation and DNA repair (72). The UV-induced nucleoid compaction observed in the BW25113 wild type (Fig. 10) was only moderate compared to previous findings in an AB1157 wild type (19) and CIP-induced compaction (Fig. 4). Nonetheless, the *ΔrecR* strain, along with *recN* and *recA* deletion strains, showed significantly impaired compaction following UV irradiation, while *ΔrecF* demonstrated only a minor delay in compaction progression compared to wild type (Fig. 10A). Importantly, these compaction impairments in the *ΔrecF* and *ΔrecR* strains did however not translate to significantly heightened UV sensitivity (Fig. 11A). The *ΔrecN* strain maintained UV sensitivity similar to wild type across varying UV doses (Fig. 11C), contrasting sharply with its remarkable sensitivity to CIP exposure (18). Combined with the short duration of UV-induced compaction (19), these findings indicate a reduced necessity for RecN-orchestrated DNA supercompaction after UV irradiation compared to after severe CIP-induced damage and strengthen our previous finding that DNA supercompaction progression depends on DNA damage severity (18).

Intriguingly, this study revealed that DNA supercompaction is prevalent across a diverse range of clinical gram-negative species isolated from various origins (Fig. 6), reinforcing our hypothesis that such compaction represents a universal cellular response to severe DNA damage. Notably, the vulnerability of the supercompaction process to disruptions from hit gene deletions was greater in the clinical reference strain CCUG17620 than in more lab-adapted strains (Fig. 5). The heightened relevance of these hit genes in clinical *E. coli* strains suggests that their role in DNA supercompaction is potentially more important within clinical settings. With their involvement in this cellular response to severe DNA damage, these genes may also influence the development of antibiotic resistance. Ultimately, continued exploration of DNA supercompaction and its underlying mechanisms is essential to evaluate its potential as a future target for developing drugs that can combat bacterial infections or prevent resistance development.

## DATA AVAILABILITY

All images from high-content imaging and the analysis results from the screening are available in the BioImage Archive (91) under accession number S-BIAD2152 at https://doi.org/10.6019/S-BIAD2152. Other data underlying this article will be shared on reasonable request to the corresponding author. All pipelines, scripts, and reference documents used for the machine learning-assisted image analysis in this study are available in the Zenodo repository at https://doi.org/10.5281/zenodo.15866742. This Zenodo repository also contains scripts for analyzing classification parameter importance, and scripts and data used for statistical analyses in R.

## SUPPLEMENTARY DATA

Supplementary Data and Supplementary Material are available online at bioRxiv.

## AUTHOR CONTRIBUTIONS

Conceptualization: K.V., N.B., K.S., E.H., J.A.B.; Methodology: K.V., N.B., I.M.M.R., S.B.R., J.V.B., K.S.; Investigation: K.V., N.B., I.M.M.R., S.B.R.; Formal analysis: K.V., N.B.; Visualization: K.V.; Software: K.V.; Funding acquisition: K.V., M.B., K.S., E.H., J.A.B.; Supervision: K.S., N.B., E.H., J.A.B.; Writing—original draft: K.V., J.A.B.; Writing—review & editing: K.V., N.B., J.V.B., K.S., E.H., J.A.B.

## ACKNOWLEDGEMENTS

We thank Anne Wahl (formerly Oslo University Hospital) for contributions to development of the high-throughput screening methodology, Stig Ove Bøe and Jon Kristen Lærdahl (Oslo University Hospital) for assistance with the high-content imaging setup and initial CellProfiler pipeline development, respectively, and Tekle Airgecho Lobie (Oslo University Hospital) for help with the SOS response assay. We are also grateful to Steven Sandler (University of Massachusetts at Amherst) for the SS6282 strain carrying HU-mCherry and Takashi Hishida (Gakushuin University) for the GFP-RecN plasmid (pSOS). Microscopy was conducted at the Advanced Light Microscopy core facility, Gaustad node, Oslo University Hospital. Screening image analyses were performed on resources provided by Sigma2 – the National Infrastructure for High-Performance Computing and Data Storage in Norway (projects nn5014k and nn9383k), as well as resources from the high-performance computing infrastructure at the University of Oslo (project ec100). Additionally, we used the NeLS and SBI portal provided by ELIXIR Norway, supported by the Research Council of Norway’s grant [270068|208481|322392] for safe storage and sharing of data from the high-content imaging. The graphical abstract was created using BioRender (https://BioRender.com/hnhuzuj). We acknowledge the use of GPT UiO, a local implementation of ChatGPT by OpenAI, for suggestions on phrasing and script code; all editing was performed by human authors.

