## Supplementary Data for "Exploring the genetic landscape of ciprofloxacin-induced DNA supercompaction in *Escherichia coli*"

**Table S1.** All 65 in-house strains included in the screening. Relevant antibiotic resistances are shown in brackets: Cam$^R$, Chloramphenicol resistance; Kan$^R$, Kanamycin resistance; Tet$^R$, Tetracycline resistance. [a] The SMG3 and ALO1208 strains are variants of the MG1655 wildtype. [b] The *recA306* notation refers to the *Δ(srl-recA)306* mutation, which only leaves a small portion of the *recA* gene and can be considered a deletion of *recA* (1–3).

| Strain | Genotype | Source |
|---|---|---|
| JABXII-72 | BW25113 ***ΔagrB*::*Kan*** [Kan$^R$] | In-house strain; deletion from (4) |
| SF169 | UF301 ***asnB3057*::*Tet*** [Tet$^R$] | In-house strain; (5, 6) |
| JAB003-F8 | BW25113 ***ΔatpBE*::*Kan*** [Kan$^R$] | In-house strain |
| CAG12077 / CGSC#7347 | MG1655 ***crcA* (*pagP*)::*Tn10*** [Tet$^R$] | (7, 8) |
| MYU001 | MG1655 ***crfC*::*Kan*** [Kan$^R$] | (9) |
| GM2927 | AB1157 ***dam-13*::*Tn9*** [Cam$^R$] | (10) |
| GM3819 | AB1157 ***dam-16*** [Kan$^R$] | (11) |
| YYH607 | YYH605 ***ΔdatA*::*Kan*** [Kan$^R$] | In-house strain; (12) |
| NL40 | DS941 ***Δdif*** [Kan$^R$] | (13) |
| JAB003-B8 | BW25113 ***ΔdinQ*::*Kan*** [Kan$^R$] | In-house strain; deletion from (4) |
| BM751 | CM735 ***dnaA204 clpX1*::*Kan*** [Kan$^R$] | (14) |
| BM750 | CM735 ***dnaA204 lon*::*Tet*** [Tet$^R$] | (14) |
| SMG379 | MG1655 ***$dnaA_{A345S}$*::*Tn10*** [Tet$^R$] | (15) |

| Strain | Genotype | Source |
|---|---|---|
| KS1117 | SMG3[a] ***dnaA*** $_{N346D}$ [Cam$^R$] | In-house strain; mutation from (16) |
| SS1512 | MG1655 ***dnaC809 zji202::Tn10*** [Tet$^R$] | In-house strain; (17) |
| JAB003-G8 | BW25113 ***dnaN*** $_{G157C}$***::Cam*** [Cam$^R$] | In-house strain; mutation from (15) |
| JAB004-A1 | BW25113 ***dnaN*** $_{G157C}$***::Cam agrB::Kan*** [Cam$^R$, Kan$^R$] | In-house strain; mutations from (4, 15) |
| SMG380 | MG1655 ***dnaN*** $_{G157C}$***::Cam*** [Cam$^R$] | (15) |
| JAB003-G3 | BW25113 ***dnaN*** $_{H175Y}$***::Tet*** [Tet$^R$] | In-house strain |
| KS1114 | SMG3[a] ***dnaN*** $_{Q156S}$ [Cam$^R$] | In-house strain; mutation from (16) |
| HI1733 | SC1148 ***dpiA::Kan*** [Kan$^R$] | (18) |
| HI1734 | SC1148 ***dpiB::Kan*** [Kan$^R$] | (18) |
| WM2016 | CSH26 ***fis::Kan*** [Kan$^R$] | (19) |
| LZ1608 | C600 ***gyrA*** $_{S83L}$ [Tet$^R$] | (20) |
| LZ3099 | C600 ***gyrA*** $_{S83L,\ D87Y}$ [Tet$^R$] | (21, 22) |
| KS1115 | SMG3[a] ***hda*** $_{F85V}$ [Tet$^R$] | In-house strain; mutation from (16) |
| ALO1387 | ALO1208[a] ***himD (ihfB)::cat*** [Cam$^R$] | In-house strain; (23) |
| ALO1415 | ALO1208[a] ***hupA16::Kan*** [Kan$^R$] | In-house strain; (23) |
| JAB003-E9 | BW25113 ***ΔibsA::Kan*** [Kan$^R$] | In-house strain |
| JAB003-E10 | BW25113 ***ΔibsB::Kan*** [Kan$^R$] | In-house strain |
| JAB003-E11 | BW25113 ***ΔibsC::Kan*** [Kan$^R$] | In-house strain |
| JAB003-E12 | BW25113 ***ΔibsD::Kan*** [Kan$^R$] | In-house strain |
| JAB003-F1 | BW25113 ***ΔibsE::Kan*** [Kan$^R$] | In-house strain |
| JAB003-B7 | BW25113 ***ΔistR::Kan*** [Kan$^R$] | In-house strain |
| JAB003-F4 | BW25113 ***ΔldrD::Kan*** [Kan$^R$] | In-house strain |
| CAG18475 | MG1655 ***metC162::Tn10*** [Tet$^R$] | (7) |
| CM735*ΔmukB* | CM735 ***ΔmukB::Kan*** [Kan$^R$] | (24) |
| FR680 | MG1655 ***mutD5 (dnaQ) zae13::Tn10*** [Tet$^R$] | (25) |
| GM7552 | GM7330 ***ΔmutH461::cat*** [Cam$^R$] | (26) |
| ME125 | MG1655 ***mutH::Kan mutL218::Tn10*** pBAD24-***MutH*** $_{E56A}$ | (27) |

| Strain | Genotype | Source |
|---|---|---|
| | P*lac-yfp*-***mutL-cat*** [Kan$^R$, Tet$^R$, Cam$^R$] | |
| KM52 | AB1157 ***mutL460*::*Cam*** [Cam$^R$] | (28) |
| ES1301 | ***mutS201*::*Tn5*** [Kan$^R$] | (29) |
| STL7742 | MG1655 ***obgE*::*Tn5*** [Kan$^R$] | (2) |
| JAB003-F3 | BW25113 ***ΔohsC*::*Kan*** (#1) [Kan$^R$] | In-house strain |
| JABXII-74 | BW25113 ***ΔohsC*::*Kan*** (#2) [Kan$^R$] | In-house strain |
| AQ7677 | MC1061 ***priA1*::*Kan*** [Kan$^R$] | (30) |
| TWx8 | MG1655 ***pyrB*::*Tn5*** [Kan$^R$] | In-house strain; (31) |
| JAB003-F5 | BW25113 ***ΔrdlD*::*Kan*** [Kan$^R$] | In-house strain |
| STL9253 | MG1655 ***recA306^b^*::*Tn10*** [Tet$^R$] | In-house strain; (1–3) |
| ALS972 | MG1655 ***recA938*::*Cam*** [Cam$^R$] | (32) |
| SMP6-346 | MG1655 ***recF*::*Tn5*** [Kan$^R$] | In-house strain |
| SS1211 | MG1655 ***Δrep*::*Cam*** [Cam$^R$] | (5) |
| IF61 | MOR166 ***ΔsdiA762*::*Kan*** [Kan$^R$] | In-house strain; mutation from (33, 34) |
| YYH609 | YYH605 ***ΔseqA*::*Tet*** [Tet$^R$] | In-house strain; (12) |
| SS211 | MG1655 pBIP-Kan ***seqAΔ10*** [Kan$^R$] | (35) |
| UF340 | AB1157 ***seqA2 asnB3057*::*Tn10*** [Tet$^R$] | (6, 36) |
| SF122 | CM735 ***seqA4 asnB3057*::*Tn10*** [Tet$^R$] | In-house strain; (6) |
| JAB003-F2 | BW25113 ***ΔshoB*::*Kan*** (#1) [Kan$^R$] | In-house strain |
| JABXII-73 | BW25113 ***ΔshoB*::*Kan*** (#2) [Kan$^R$] | In-house strain |
| JAB003-F7 | BW25113 ***ΔspoT*::*Kan*** [Kan$^R$] | In-house strain; deletion from (34) |
| JAB003-B6 | BW25113 ***ΔtisB*::*Kan*** [Kan$^R$] | In-house strain |
| LBB925 | LBB451 ***tolC*::*Tn10*** [Tet$^R$] | In-house strain |
| MG1655-C122 | MG1655 ***umuC122*::*Tn5*** [Kan$^R$] | In-house strain; (37) |
| DS984 | DS941 ***xerC*::*Mu*** [Cam$^R$] | (13, 38) |
| DS9008 | DS941 ***xerD*::*Tn10*** [Tet$^R$] | (13, 39) |

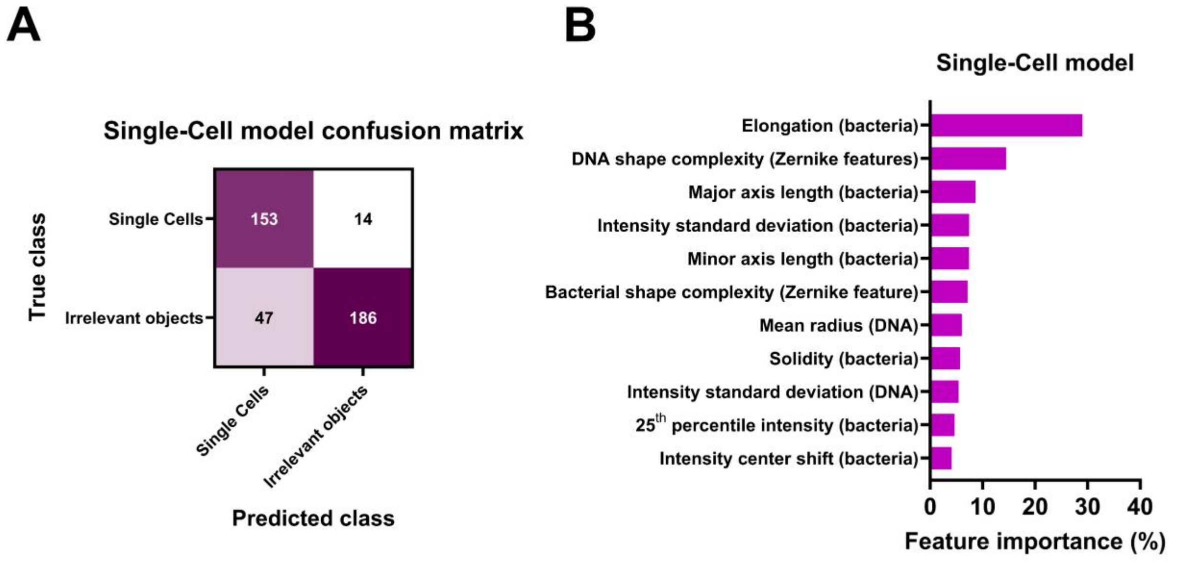


**Figure S1.** The Single-Cell model was trained to distinguish single cells from irrelevant objects. (**A**) Confusion matrix for the Single-Cell model quantifying its prediction performance. The matrix displays the true phenotypes for 200 cells per model-predicted class. (**B**) Feature importance quantification for the Single-Cell model, illustrating contribution of various parameters to classification of single cells. Parameters are grouped based on similarity to emphasize which cellular features are critical for accurate detection of single cells.

**Table S2.** Parameters used in the Single-Cell model, defining criteria for classifying single cells versus irrelevant objects. Each parameter's associated CellProfiler module and measurement are described, along with interpretations of the assessed cellular features. All measurements were performed on segmented objects (cells). Classification weights are provided for single cell (S) and irrelevant object (I) classes, based on measurements above (true) and below (false) threshold values. Thresholds are rounded to four decimals; weights to two decimals.

| Rule # | CellProfiler module and measurement | Feature interpretation | Threshold | Weight (true) | Weight (false) |
|---|---|---|---|---|---|
| 1 | AreaShape-Eccentricity | Elongation (bacteria) | 0.8676 | S: +0.50<br>I: -0.50 | S: -0.86<br>I: +0.86 |
| 2 | RadialDistribution-ZernikeMagnitude-EnhancedDNAImage-2-0 | DNA shape complexity (Zernike features) | 0.0022 | S: +0.26<br>I: -0.26 | S: -0.93<br>I: +0.93 |
| 3 | Mean-DNAObjects-AreaShape-MeanRadius | Mean radius (DNA) | 1.7241 | S: +0.17<br>I: -0.17 | S: -0.56<br>I: +0.56 |
| 4 | AreaShape-MinorAxisLength | Minor axis length (bacteria) | 14.0563 | S: -0.71<br>I: +0.71 | S: +0.19<br>I: -0.19 |
| 5 | AreaShape-MajorAxisLength | Major axis length (bacteria) | 17.6847 | S: +0.09<br>I: -0.09 | S: -0.95<br>I: +0.95 |
| 6 | AreaShape-Eccentricity | Elongation (bacteria) | 0.9306 | S: +0.29<br>I: -0.29 | S: -0.27<br>I: +0.27 |
| 7 | AreaShape-Solidity | Solidity (bacteria) | 0.8298 | S: +0.33<br>I: -0.33 | S: -0.36<br>I: +0.36 |
| 8 | Intensity-StdIntensity-RawBacterialmage | Intensity standard deviation (bacteria) | 0.0004 | S: +0.09<br>I: -0.09 | S: -0.81<br>I: +0.81 |
| 9 | Mean-DNAObjects-AreaShape-Zernike-2-0 | DNA shape complexity (Zernike features) | 0.1033 | S: +0.15<br>I: -0.15 | S: -0.40<br>I: +0.40 |
| 10 | AreaShape-Eccentricity | Elongation (bacteria) | 0.8103 | S: +0.07<br>I: -0.07 | S: -0.90<br>I: +0.90 |
| 11 | AreaShape-Zernike-3-1 | Bacterial shape complexity (Zernike feature) | 0.0045 | S: -0.09<br>I: +0.09 | S: +0.77<br>I: -0.77 |
| 12 | Intensity-MassDisplacement-RawBacterialmage | Intensity center shift (bacteria) | 0.2507 | S: -0.35<br>I: +0.35 | S: +0.14<br>I: -0.14 |
| 13 | AreaShape-Eccentricity | Elongation (bacteria) | 0.9555 | S: +0.46<br>I: -0.46 | S: -0.14<br>I: +0.14 |

| Rule # | CellProfiler module and measurement | Feature interpretation | Threshold | Weight (true) | Weight (false) |
|---|---|---|---|---|---|
| 14 | Intensity-LowerQuartileIntensity-EnhancedBacteriaimage | 25th percentile intensity (bacteria) | 0.0121 | S: +0.42<br>I: -0.42 | S: -0.14<br>I: +0.14 |
| 15 | Intensity-StdIntensity-DNAImage | Intensity standard deviation (DNA) | 0.0036 | S: +0.07<br>I: -0.07 | S: -0.58<br>I: +0.58 |

**Table S3.** Parameters used in the Phenotype model, defining criteria for classifying the different DNA compaction phenotypes. Each parameter's associated CellProfiler module and measurement are described, along with interpretations of the assessed cellular features. All measurements were performed on single cells identified by the Single-Cell model. Classification weights are provided for the wild-type (W), unchallenged (U) and ΔrecN (R) phenotypes, based on measurements above (true) and below (false) threshold value. Thresholds are rounded to four decimals; weights to two decimals.

| Rule # | CellProfiler module and measurement | Feature interpretation | Threshold | Weight (true) | Weight (false) |
|---|---|---|---|---|---|
| 1 | RadialDistribution-MeanFrac-EnhancedDNAImage-1of6 | Mean midcell intensity (DNA) | 2.9628 | W: +0.66<br>U: -0.96<br>R: -0.70 | W: -0.79<br>U: -0.05<br>R: -0.16 |
| 2 | RadialDistribution-MeanFrac-EnhancedDNAImage-1of6 | Mean midcell intensity (DNA) | 2.0326 | W: +0.00<br>U: -0.57<br>R: +0.18 | W: -0.46<br>U: +0.49<br>R: -0.48 |
| 3 | Children-SingleCellDNAObjects-Count | DNA foci count | 1 | W: -0.43<br>U: +0.05<br>R: +0.25 | W: +0.51<br>U: -0.20<br>R: -0.48 |
| 4 | Intensity-StdIntensity-SingleCellDNAImage | Intensity standard deviation (DNA) | 0.0084 | W: +0.48<br>U: -0.29<br>R: -0.21 | W: -0.43<br>U: +0.13<br>R: +0.08 |
| 5 | Mean-SingleCellDNAObjects-AreaShape-MinorAxisLength | Mean minor axis length (DNA) | 6.0858 | W: -0.20<br>U: +0.06<br>R: +0.01 | W: +0.83<br>U: -0.59<br>R: -0.11 |
| 6 | Intensity-MedianIntensity-EnhancedDNAImage | Median intensity (DNA) | 0.0104 | W: -0.64<br>U: +0.37<br>R: -0.14 | W: +0.20<br>U: -0.19<br>R: +0.05 |
| 7 | RadialDistribution-ZernikeMagnitude-EnhancedDNAImage-8-4 | DNA shape complexity (Zernike features) | 0.0003 | W: -0.36<br>U: -0.01<br>R: +0.09 | W: +0.32<br>U: +0.04<br>R: -0.17 |
| 8 | RadialDistribution-RadialCV-RawDNAImage-1of6 | Midcell intensity variability (DNA) | 0.0614 | W: +0.13<br>U: -0.30<br>R: +0.18 | W: -0.23<br>U: +0.25<br>R: -0.18 |
| 9 | RadialDistribution-ZernikeMagnitude-SingleCellDNAImage-3-1 | DNA shape complexity (Zernike features) | 0.0028 | W: +0.36<br>U: +0.02<br>R: -0.17 | W: -0.31<br>U: -0.01<br>R: +0.08 |
| 10 | RadialDistribution-MeanFrac-RawDNAImage-1of6 | Mean midcell intensity (DNA) | 1.4091 | W: -0.08<br>U: -0.07<br>R: +0.10 | W: +0.46<br>U: +0.39<br>R: -0.56 |
| 11 | RadialDistribution-ZernikeMagnitude-EnhancedDNAImage- | DNA shape complexity | 0.0002 | W: -0.12<br>U: +0.01 | W: +0.65<br>U: -0.11 |

| Rule # | CellProfiler module and measurement | Feature interpretation | Threshold | Weight (true) | Weight (false) |
|---|---|---|---|---|---|
| | 5-3 | (Zernike features) | | R: +0.05 | R: -0.48 |
| 12 | RadialDistribution-MeanFrac-RawDNAImage-1of6 | Mean midcell intensity (DNA) | 3.1113 | W: +0.89<br>U: -1.00<br>R: -0.85 | W: -0.04<br>U: +0.02<br>R: +0.03 |
| 13 | RadialDistribution-ZernikeMagnitude-SingleCellDNAImage-9-5 | DNA shape complexity (Zernike features) | 0.0008 | W: -0.60<br>U: +0.14<br>R: +0.09 | W: +0.14<br>U: -0.04<br>R: -0.03 |
| 14 | Intensity-StdIntensity-SingleCellDNAImage | Intensity standard deviation (DNA) | 0.0064 | W: +0.23<br>U: -0.11<br>R: -0.02 | W: -0.39<br>U: +0.10<br>R: +0.02 |
| 15 | Mean-SingleCellDNAObjects-AreaShape-Zernike-2-2 | DNA shape complexity (Zernike features) | 0.1138 | W: +0.39<br>U: -0.34<br>R: +0.13 | W: -0.16<br>U: +0.07<br>R: -0.03 |

**Table S4.** The 156 initial candidate strains from the screening. [a] The *recA306* notation refers to the *Δ(srl-recA)306* mutation, which only leaves a small portion of the *recA* gene and can be considered a deletion of *recA* (1–3).

| Strain | Genotype | Source |
|---|---|---|
| JW3713 | BW25113 ***ΔatpH*** | Keio collection (34) |
| JW0865 | BW25113 ***ΔclpS*** | Keio collection |
| JW1241 | BW25113 ***Δcls* (*clsA*)** | Keio collection |
| JW1106 | BW25113 ***ΔcobB*** | Keio collection |
| JW2406 | BW25113 ***ΔcysZ*** | Keio collection |
| GM3819 | AB1157 ***dam-16*** | In-house strain; (11) |
| JW3351 | BW25113 ***ΔdamX*** | Keio collection |
| JW0613 | BW25113 ***ΔdcuC*** | Keio collection |
| JW0090 | BW25113 ***ΔddlB*** | Keio collection |
| JW3670 | BW25113 ***ΔdgoK*** | Keio collection |
| JW5187 | BW25113 ***ΔdhaK*** | Keio collection |
| JW0054 | BW25113 ***ΔdjlA*** | Keio collection |
| JW3228 | BW25113 ***ΔdusB*** | Keio collection |
| JW2661 | BW25113 ***ΔemrB*** | Keio collection |
| JW1077 | BW25113 ***ΔfabH*** | Keio collection |
| JW5020 | BW25113 ***ΔfadE*** | Keio collection |
| JW4191 | BW25113 ***Δfbp*** | Keio collection |
| JW1379 | BW25113 ***ΔfeaR*** | Keio collection |
| JW0149 | BW25113 ***ΔfhuB*** | Keio collection |
| WM2016 | CSH26 ***fis*::*Kan*** | In-house strain; (19) |
| JW1058 | BW25113 ***ΔflgM*** | Keio collection |
| JW1907 | BW25113 ***ΔfliA*** | Keio collection |
| JW4112 | BW25113 ***ΔfrdD*** | Keio collection |
| JW3869 | BW25113 ***ΔfrvX*** | Keio collection |
| JW1413 | BW25113 ***ΔgapC*** | Keio collection |
| JW3841 | BW25113 ***ΔglnA*** | Keio collection |
| JW3896 | BW25113 ***ΔglpX*** | Keio collection |
| JW3530 | BW25113 ***ΔglyS*** | Keio collection |
| JW3297 | BW25113 ***ΔgspO*** | Keio collection |
| JW0173 | BW25113 ***ΔhlpA* (*skp*)** | Keio collection |
| JW2965 | BW25113 ***ΔhybO*** | Keio collection |

| **Strain** | **Genotype** | **Source** |
|---|---|---|
| JW2470 | BW25113 ***ΔhyfE*** | Keio collection |
| JW4221 | BW25113 ***ΔidnR*** | Keio collection |
| JW0076 | BW25113 ***ΔilvI*** | Keio collection |
| JW0687 | BW25113 ***ΔkdpF*** | Keio collection |
| JW0071 | BW25113 ***ΔleuC*** | Keio collection |
| JW0429 | BW25113 ***Δlon*** | Keio collection |
| JW1041 | BW25113 ***ΔlpxL*** | Keio collection |
| JW1145 | BW25113 ***ΔmcrA*** | Keio collection |
| JW5338 | BW25113 ***ΔmdtA*** | Keio collection |
| JW0341 | BW25113 ***ΔmhpD*** | Keio collection |
| CM735*ΔmukB* | CM735 ***ΔmukB*::*Kan*** | In-house strain; (24) |
| JW4128 | BW25113 ***ΔmutL*** | Keio collection |
| JW3002 | BW25113 ***ΔnudF*** | Keio collection |
| JW5375 | BW25113 ***ΔnuoC*** | Keio collection |
| JW2280 | BW25113 ***ΔnuoE*** | Keio collection |
| JW2279 | BW25113 ***ΔnuoF*** | Keio collection |
| JW2278 | BW25113 ***ΔnuoG*** | Keio collection |
| JW2277 | BW25113 ***ΔnuoH*** | Keio collection |
| STL7742 | MG1655 ***obgE*::*Tn5*** | In-house strain; (2) |
| JW0912 | BW25113 ***ΔompF*** | Keio collection |
| JW1113 | BW25113 ***ΔpepT*** | Keio collection |
| JW3985 | BW25113 ***Δpgi*** | Keio collection |
| JW1110 | BW25113 ***ΔpotC*** | Keio collection |
| JW0104 | BW25113 ***ΔppdD*** | Keio collection |
| STL9253 | MG1655 ***recA306*[a]::*Tn10*** | In-house strain; (1–3) |
| ALS972 | MG1655 ***recA938*::*Cam*** | In-house strain; (32) |
| JW2788 | BW25113 ***ΔrecB*** | Keio collection |
| JW2790 | BW25113 ***ΔrecC*** | Keio collection |
| JW2787 | BW25113 ***ΔrecD*** | Keio collection |
| JW3677 | BW25113 ***ΔrecF*** | Keio collection |
| JW5855 | BW25113 ***ΔrecQ*** | Keio collection |
| JW0461 | BW25113 ***ΔrecR*** | Keio collection |
| JW3946 | BW25113 ***ΔrplK*** | Keio collection |
| JW3907 | BW25113 ***ΔrpmE*** | Keio collection |

| Strain | Genotype | Source |
|---|---|---|
| JW1707 | BW25113 ***ΔrpmI*** | Keio collection |
| JW3039 | BW25113 ***ΔrpoD*** | Keio collection |
| JW3169 | BW25113 ***ΔrpoN*** | Keio collection |
| JW0387 | BW25113 ***ΔsbcC*** | Keio collection |
| JW0388 | BW25113 ***ΔsbcD*** | Keio collection |
| JW0711 | BW25113 ***ΔsdhC*** | Keio collection |
| JAB003-F2 | BW25113 ***ΔshoB::Kan*** (#1) | In-house strain |
| JW3245 | BW25113 ***Δsmg*** | Keio collection |
| JW1264 | BW25113 ***ΔsohB*** | Keio collection |
| JW5404 | BW25113 ***ΔsseB*** | Keio collection |
| JW5273 | BW25113 ***ΔsufB*** | Keio collection |
| JW2985 | BW25113 ***ΔsufI*** (***ftsP***) | Keio collection |
| JW1425 | BW25113 ***ΔtehA*** | Keio collection |
| JW0979 | BW25113 ***ΔtorT*** | Keio collection |
| JW4019 | BW25113 ***ΔuvrA*** | Keio collection |
| JW5892 | BW25113 ***ΔyadB*** (***gluQ***) | Keio collection |
| JW0349 | BW25113 ***ΔyaiO*** | Keio collection |
| JW0362 | BW25113 ***ΔyaiT*** | Keio collection |
| JW0366 | BW25113 ***ΔyaiV*** (***iprA***) | Keio collection |
| JW0369 | BW25113 ***ΔyaiW*** | Keio collection |
| JW0370 | BW25113 ***ΔyaiY*** | Keio collection |
| JW0492 | BW25113 ***ΔybbS*** (***allS***) | Keio collection |
| JW0527 | BW25113 ***ΔybcD*** (***peaD***) | Keio collection |
| JW0532 | BW25113 ***ΔybcK*** | Keio collection |
| JW0702 | BW25113 ***ΔybgK*** (***pxpC***) | Keio collection |
| JW5106 | BW25113 ***ΔybiM*** (***mcbA***) | Keio collection |
| JW0910 | BW25113 ***ΔycbL*** (***gloC***) | Keio collection |
| JW0924 | BW25113 ***ΔycbT*** (***elfG***) | Keio collection |
| JW0942 | BW25113 ***ΔyccR (sxy)*** | Keio collection |
| JW5142 | BW25113 ***ΔycdR*** (***pgaB***) | Keio collection |
| JW1042 | BW25113 ***ΔyceA (trhO)*** | Keio collection |
| JW1074 | BW25113 ***ΔyceD*** | Keio collection |
| JW1246 | BW25113 ***ΔyciB*** | Keio collection |
| JW5196 | BW25113 ***ΔyciO*** | Keio collection |
| JW1313 | BW25113 ***ΔycjW*** | Keio collection |

| **Strain** | **Genotype** | **Source** |
|---|---|---|
| JW5207 | BW25113 ***ΔydaQ*** **(*xisR*)** | Keio collection |
| JW5215 | BW25113 ***ΔydbJ*** | Keio collection |
| JW5228 | BW25113 ***ΔydcM*** **(*insQ*)** | Keio collection |
| JW1459 | BW25113 ***ΔyddE*** | Keio collection |
| JW1527 | BW25113 ***ΔydeE*** | Keio collection |
| JW1530 | BW25113 ***ΔydeJ*** | Keio collection |
| JW5243 | BW25113 ***ΔydeN*** | Keio collection |
| JW1637 | BW25113 ***ΔydhK*** | Keio collection |
| JW1659 | BW25113 ***ΔydhT*** | Keio collection |
| JW5271 | BW25113 ***ΔydhX*** | Keio collection |
| JW5274 | BW25113 ***ΔydiN*** | Keio collection |
| JW5293 | BW25113 ***ΔyeaV*** | Keio collection |
| JW5313 | BW25113 ***ΔyedO*** **(*dcyD*)** | Keio collection |
| JW2088 | BW25113 ***ΔyegW*** | Keio collection |
| JW2400 | BW25113 ***ΔyfeR*** | Keio collection |
| JW2421 | BW25113 ***ΔyfeU*** **(*murQ*)** | Keio collection |
| JW5458 | BW25113 ***ΔygeK*** | Keio collection |
| JW3040 | BW25113 ***ΔygjF*** **(*mug*)** | Keio collection |
| JW3057 | BW25113 ***ΔygjQ*** | Keio collection |
| JW3092 | BW25113 ***ΔyhaC*** | Keio collection |
| JW3125 | BW25113 ***ΔyhbS*** | Keio collection |
| JW3129 | BW25113 ***ΔyhbW*** | Keio collection |
| JW3255 | BW25113 ***ΔyhdN*** | Keio collection |
| JW3454 | BW25113 ***ΔyhiI*** | Keio collection |
| JW3557 | BW25113 ***ΔyiaU*** | Keio collection |
| JW3631 | BW25113 ***ΔyicI*** | Keio collection |
| JW3689 | BW25113 ***ΔyidZ*** | Keio collection |
| JW3831 | BW25113 ***ΔyihE*** **(*srkA*)** | Keio collection |
| JW5568 | BW25113 ***ΔyihV*** | Keio collection |
| JW3899 | BW25113 ***ΔyiiU*** **(*zapB*)** | Keio collection |
| JW3936 | BW25113 ***ΔyijD*** | Keio collection |
| JW3971 | BW25113 ***ΔyjaA*** | Keio collection |
| JW3989 | BW25113 ***ΔyjbH*** | Keio collection |
| JW4017 | BW25113 ***ΔyjbQ*** | Keio collection |
| JW4018 | BW25113 ***ΔyjbR*** | Keio collection |

| **Strain** | **Genotype** | **Source** |
|---|---|---|
| JW4026 | BW25113 ***ΔyjcE*** | Keio collection |
| JW4042 | BW25113 ***ΔyjcQ* (*mdtO*)** | Keio collection |
| JW4105 | BW25113 ***ΔyjeJ*** | Keio collection |
| JW4124 | BW25113 ***ΔyjeS* (*queG*)** | Keio collection |
| JW4148 | BW25113 ***ΔyjfP*** | Keio collection |
| JW4271 | BW25113 ***ΔyjhR*** | Keio collection |
| JW5968 | BW25113 ***ΔyjhX* (*topAI*)** | Keio collection |
| JW4354 | BW25113 ***ΔyjjK* (*ettA*)** | Keio collection |
| JW4365 | BW25113 ***ΔyjjY*** | Keio collection |
| JW5039 | BW25113 ***ΔykgI* (*rclB*)** | Keio collection |
| JW5133 | BW25113 ***ΔymcD* (*gfcA*)** | Keio collection |
| JW1154 | BW25113 ***ΔymgC*** | Keio collection |
| JW5230 | BW25113 ***ΔyncN* (*hicA*)** | Keio collection |
| JW5244 | BW25113 ***ΔyneL*** | Keio collection |
| JW5251 | BW25113 ***ΔynfO*** | Keio collection |
| JW5331 | BW25113 ***ΔyoeB*** | Keio collection |
| JW2460 | BW25113 ***ΔypfJ*** | Keio collection |
| JW3111 | BW25113 ***ΔyraH*** | Keio collection |
| JW3356 | BW25113 ***ΔyrfB* (*hofO*)** | Keio collection |
| JW4167 | BW25113 ***ΔytfE*** | Keio collection |
| JW4169 | BW25113 ***ΔytfG* (*qorB*)** | Keio collection |

**Table S5.** All 93 poorly growing strains from the screening that were re-imaged. [a] The SMG3 and ALO1208 strains are variants of the MG1655 wildtype.

| Strain | Genotype | Source |
|---|---|---|
| JW3234 | BW25113 ***ΔacrF*** | Keio collection (34) |
| JW2201 | BW25113 ***Δada*** | Keio collection |
| JW1228 | BW25113 ***ΔadhE*** | Keio collection |
| JW4049 | BW25113 ***ΔalsB*** | Keio collection |
| JW4047 | BW25113 ***ΔalsC*** | Keio collection |
| JW4046 | BW25113 ***ΔalsE*** | Keio collection |
| JW4111 | BW25113 ***ΔampC*** | Keio collection |
| JW3470 | BW25113 ***ΔarsC*** | Keio collection |
| JW5239 | BW25113 ***Δbdm*** | Keio collection |
| JW0305 | BW25113 ***ΔbetI*** | Keio collection |
| JW2869 | BW25113 ***ΔbglA*** | Keio collection |
| CAG12077 / CGSC#7347 | MG1655 ***crcA* (*pagP*)::*Tn10*** | In-house strain; (7, 8) |
| JW2639 | BW25113 ***ΔcsiR* (*glaR*)** | Keio collection |
| JW1863 | BW25113 ***ΔcutC*** | Keio collection |
| JW2416 | BW25113 ***ΔcysW*** | Keio collection |
| JW3905 | BW25113 ***ΔcytR*** | Keio collection |
| JW5592 | BW25113 ***ΔdapF*** | Keio collection |
| BM750 | CM735 ***dnaA204 lon*::*Tet*** | In-house strain; (14) |
| SMG379 | MG1655 ***$dnaA_{A345S}$*::*Tn10*** | In-house strain; (15) |
| HI1733 | SC1148 ***dpiA*::*Kan*** | In-house strain; (18) |
| HI1734 | SC1148 ***dpiB*::*Kan*** | In-house strain; (18) |
| JW3509 | BW25113 ***ΔdppF*** | Keio collection |
| JW1506 | BW25113 ***Δego* (*lsrA*)** | Keio collection |
| JW2365 | BW25113 ***ΔemrK*** | Keio collection |
| JW2434 | BW25113 ***ΔeutB*** | Keio collection |
| JW2437 | BW25113 ***ΔeutG*** | Keio collection |
| JW2366 | BW25113 ***ΔevgA*** | Keio collection |
| JW3065 | BW25113 ***ΔexuR*** | Keio collection |
| JW4250 | BW25113 ***ΔfecB*** | Keio collection |
| JW1070 | BW25113 ***ΔflgL*** | Keio collection |
| JW1328 | BW25113 ***Δfnr*** | Keio collection |

| Strain | Genotype | Source |
|---|---|---|
| JW3333 | BW25113 ***ΔfrlA*** | Keio collection |
| LZ1608 | C600 ***gyrA**$_{S83L}$* | In-house strain; (20) |
| JW1950 | BW25113 ***ΔhchA*** | Keio collection |
| KS1115 | SMG3[a] ***hda**$_{F85V}$* | In-house strain |
| JW4130 | BW25113 ***Δhfq*** | Keio collection |
| ALO1387 | ALO1208[a] ***himD* (*ihfB*)::*cat*** | In-house strain; (23) |
| JW0645 | BW25113 ***ΔhscC*** | Keio collection |
| JW3903 | BW25113 ***ΔhslV*** | Keio collection |
| JW3903 | BW25113 ***ΔhybA*** | Keio collection |
| JW3978 | BW25113 ***ΔiclR*** | Keio collection |
| JW0895 | BW25113 ***ΔihfB*** | Keio collection |
| JW3313 | BW25113 ***ΔkefB*** | Keio collection |
| JW0334 | BW25113 ***ΔlacY*** | Keio collection |
| JW0720 | BW25113 ***ΔmngA*** | Keio collection |
| JW3573 | BW25113 ***ΔmtlA*** | Keio collection |
| FR680 | MG1655 ***mutD5* (*dnaQ*) *zae13*::*Tn10*** | In-house strain; (25) |
| JW3193 | BW25113 ***ΔnanT*** | Keio collection |
| JW2712 | BW25113 ***ΔnlpD*** | Keio collection |
| JW4033 | BW25113 ***ΔnrfC*** | Keio collection |
| JW5875 | BW25113 ***ΔnuoB*** | Keio collection |
| JW2273 | BW25113 ***ΔnuoL*** | Keio collection |
| JW4064 | BW25113 ***ΔphnE*** | Keio collection |
| JW1555 | BW25113 ***ΔrelE*** | Keio collection |
| JW3753 | BW25113 ***ΔrhlB*** | Keio collection |
| JW3756 | BW25113 ***Δrho*** | Keio collection |
| JW1072 | BW25113 ***ΔrluC*** | Keio collection |
| JW3947 | BW25113 ***ΔrplA*** | Keio collection |
| JW4161 | BW25113 ***ΔrplI*** | Keio collection |
| JW3261 | BW25113 ***ΔrpmJ*** | Keio collection |
| JW3134 | BW25113 ***ΔrpsO*** | Keio collection |
| JW2132 | BW25113 ***ΔsanA*** | Keio collection |
| JW1284 | BW25113 ***ΔsapD*** | Keio collection |
| JW5967 | BW25113 ***ΔsgcB*** | Keio collection |
| JW2598 | BW25113 ***ΔsmpA* (*bamE*)** | Keio collection |
| JW5962 | BW25113 ***Δsra*** | Keio collection |

| Strain | Genotype | Source |
|---|---|---|
| JW2847 | BW25113 ***ΔssnA*** | Keio collection |
| JW0919 | BW25113 ***ΔssuA*** | Keio collection |
| JW5738 | BW25113 ***ΔsugE* (*gdx*)** | Keio collection |
| JW0622 | BW25113 ***ΔtatE*** | Keio collection |
| JW0067 | BW25113 ***ΔtbpA* (*thiB*)** | Keio collection |
| LBB925 | LBB451 ***tolC::Tn10*** | In-house strain |
| JW4154 | BW25113 ***ΔulaD*** | Keio collection |
| JW2483 | BW25113 ***Δupp*** | Keio collection |
| JW3991 | BW25113 ***ΔxylE*** | Keio collection |
| JW0222 | BW25113 ***ΔyafN*** | Keio collection |
| JW0279 | BW25113 ***ΔyagS* (*paoB*)** | Keio collection |
| JW0445 | BW25113 ***ΔybaA*** | Keio collection |
| JW0696 | BW25113 ***ΔybfD*** | Keio collection |
| JW0725 | BW25113 ***ΔybgE*** | Keio collection |
| JW0780 | BW25113 ***ΔybiH* (*cecR*)** | Keio collection |
| JW1015 | BW25113 ***ΔycdU*** | Keio collection |
| JW1271 | BW25113 ***ΔyciS* (*lapA*)** | Keio collection |
| JW5198 | BW25113 ***ΔyciX*** | Keio collection |
| JW5252 | BW25113 ***ΔydfO*** | Keio collection |
| JW1859 | BW25113 ***ΔyecO* (*cmoA*)** | Keio collection |
| JW3253 | BW25113 ***ΔyhdL* (*arfA*)** | Keio collection |
| JW3750 | BW25113 ***ΔyifN*** | Keio collection |
| JW5756 | BW25113 ***ΔyjgK* (*tabA*)** | Keio collection |
| JW4310 | BW25113 ***ΔyjiW* (*symE*)** | Keio collection |
| JW4341 | BW25113 ***ΔyjjV*** | Keio collection |
| JW5245 | BW25113 ***ΔyneE*** | Keio collection |
| JW4168 | BW25113 ***ΔytfF*** | Keio collection |

**Table S6.** Remaining 54 candidate strains after re-imaging of initial candidates and poorly growing strains. [a] The SMG3 strain is a variant of the MG1655 wildtype. [b] The *recA306* notation refers to the *Δ(srl-recA)306* mutation, which only leaves a small portion of the *recA* gene and can be considered a deletion of *recA* (1–3).

| Strain | Genotype | Source |
|---|---|---|
| JW2201 | BW25113 ***Δada*** | Keio collection (34) |
| JW1228 | BW25113 ***ΔadhE*** | Keio collection |
| JW4047 | BW25113 ***ΔalsC*** | Keio collection |
| JW3713 | BW25113 ***ΔatpH*** | Keio collection |
| JW0865 | BW25113 ***ΔclpS*** | Keio collection |
| JW3905 | BW25113 ***ΔcytR*** | Keio collection |
| HI1733 | SC1148 ***dpiA*::*Kan*** | In-house strain; (18) |
| HI1734 | SC1148 ***dpiB*::*Kan*** | In-house strain; (18) |
| JW3509 | BW25113 ***ΔdppF*** | Keio collection |
| JW3228 | BW25113 ***ΔdusB*** | Keio collection |
| JW1506 | BW25113 ***Δego* (*lsrA*)** | Keio collection |
| JW1077 | BW25113 ***ΔfabH*** | Keio collection |
| WM2016 | CSH26 ***fis*::*Kan*** | In-house strain; (19) |
| JW1328 | BW25113 ***Δfnr*** | Keio collection |
| JW4112 | BW25113 ***ΔfrdD*** | Keio collection |
| JW1950 | BW25113 ***ΔhchA*** | Keio collection |
| KS1115 | SMG3[a] $\boldsymbol{hda_{F85V}}$ | In-house strain |
| JW4130 | BW25113 ***Δhfq*** | Keio collection |
| JW2470 | BW25113 ***ΔhyfE*** | Keio collection |
| JW0895 | BW25113 ***ΔihfB*** | Keio collection |
| JW3313 | BW25113 ***ΔkefB*** | Keio collection |
| JW1041 | BW25113 ***ΔlpxL*** | Keio collection |
| JW3573 | BW25113 ***ΔmtlA*** | Keio collection |
| JW5875 | BW25113 ***ΔnuoB*** | Keio collection |
| STL7742 | MG1655 ***obgE*::*Tn5*** | In-house strain; (2) |
| STL9253 | MG1655 ***recA306*[b]::*Tn10*** | In-house strain; (1–3) |

| Strain | Genotype | Source |
|---|---|---|
| ALS972 | MG1655 ***recA938*::*Cam*** | In-house strain; (32) |
| JW2788 | BW25113 ***ΔrecB*** | Keio collection |
| JW2790 | BW25113 ***ΔrecC*** | Keio collection |
| JW2787 | BW25113 ***ΔrecD*** | Keio collection |
| JW3677 | BW25113 ***ΔrecF*** | Keio collection |
| JW5855 | BW25113 ***ΔrecQ*** | Keio collection |
| JW0461 | BW25113 ***ΔrecR*** | Keio collection |
| JW1707 | BW25113 ***ΔrpmI*** | Keio collection |
| JW0387 | BW25113 ***ΔsbcC*** | Keio collection |
| JW0388 | BW25113 ***ΔsbcD*** | Keio collection |
| JW5404 | BW25113 ***ΔsseB*** | Keio collection |
| JW0622 | BW25113 ***ΔtatE*** | Keio collection |
| JW0067 | BW25113 ***ΔtbpA* (*thiB*)** | Keio collection |
| JW1425 | BW25113 ***ΔtehA*** | Keio collection |
| JW0979 | BW25113 ***ΔtorT*** | Keio collection |
| JW0369 | BW25113 ***ΔyaiW*** | Keio collection |
| JW0370 | BW25113 ***ΔyaiY*** | Keio collection |
| JW1527 | BW25113 ***ΔydeE*** | Keio collection |
| JW1530 | BW25113 ***ΔydeJ*** | Keio collection |
| JW3255 | BW25113 ***ΔyhdN*** | Keio collection |
| JW3899 | BW25113 ***ΔyiiU* (*zapB*)** | Keio collection |
| JW4018 | BW25113 ***ΔyjbR*** | Keio collection |
| JW4026 | BW25113 ***ΔyjcE*** | Keio collection |
| JW4042 | BW25113 ***ΔyjcQ* (*mdtO*)** | Keio collection |
| JW4310 | BW25113 ***ΔyjiW* (*symE*)** | Keio collection |
| JW4365 | BW25113 ***ΔyjjY*** | Keio collection |
| JW5244 | BW25113 ***ΔyneL*** | Keio collection |
| JW4167 | BW25113 ***ΔytfE*** | Keio collection |

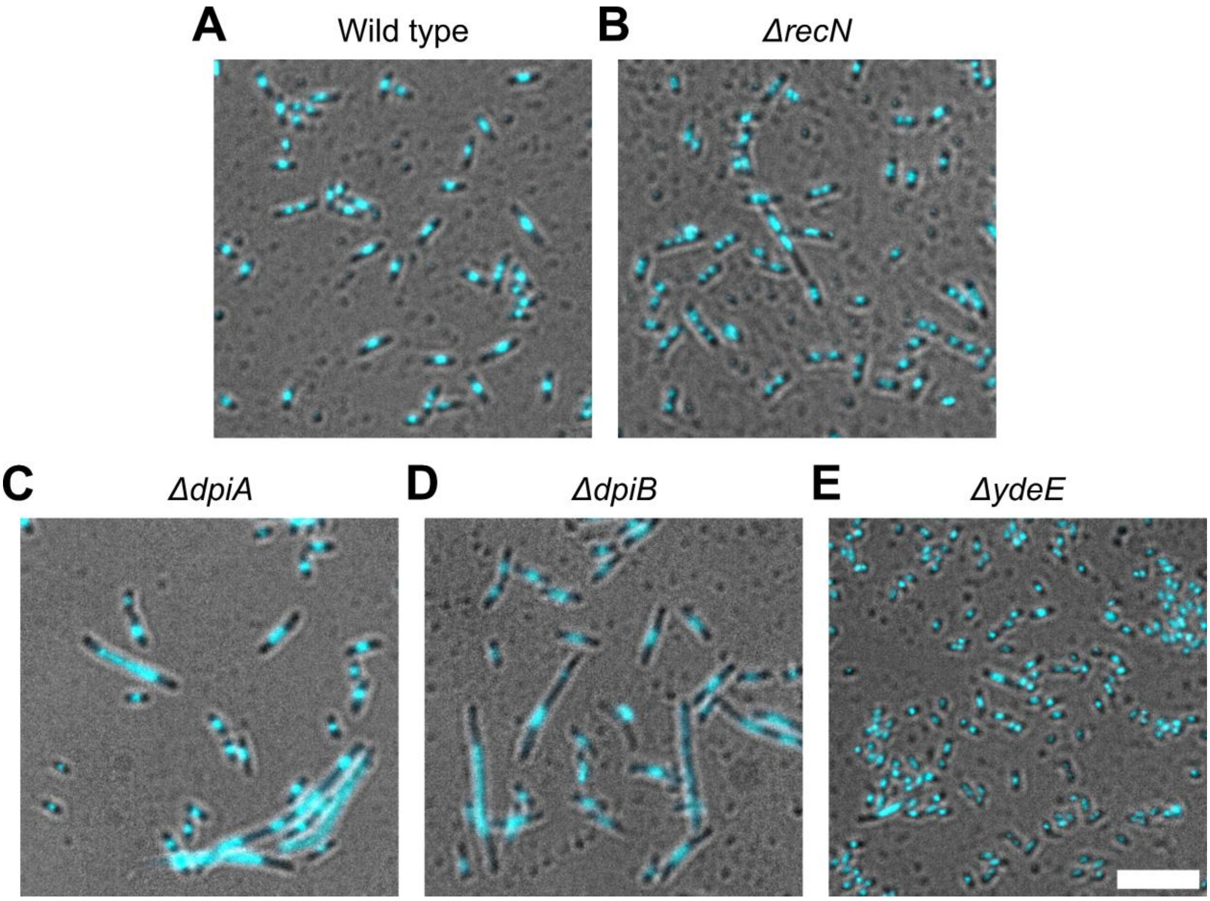


**Figure S2.** Example images of various strains with distinctive phenotypes captured using high-content imaging after CIP exposure. Strains were cultured, exposed to CIP for 15-20 minutes, and prepared for imaging in 384-well plates. (**A**) Wild type and (**B**) the ΔrecN strain served as controls in the screening. (**C**) ΔdpiA and (**D**) ΔdpiB strains only demonstrated impaired supercompaction in a subpopulation of filamenting cells. (**E**) The ΔydeE strain exhibits notably short and small cells. All images are at the same scale; scale bar is 10 µm.

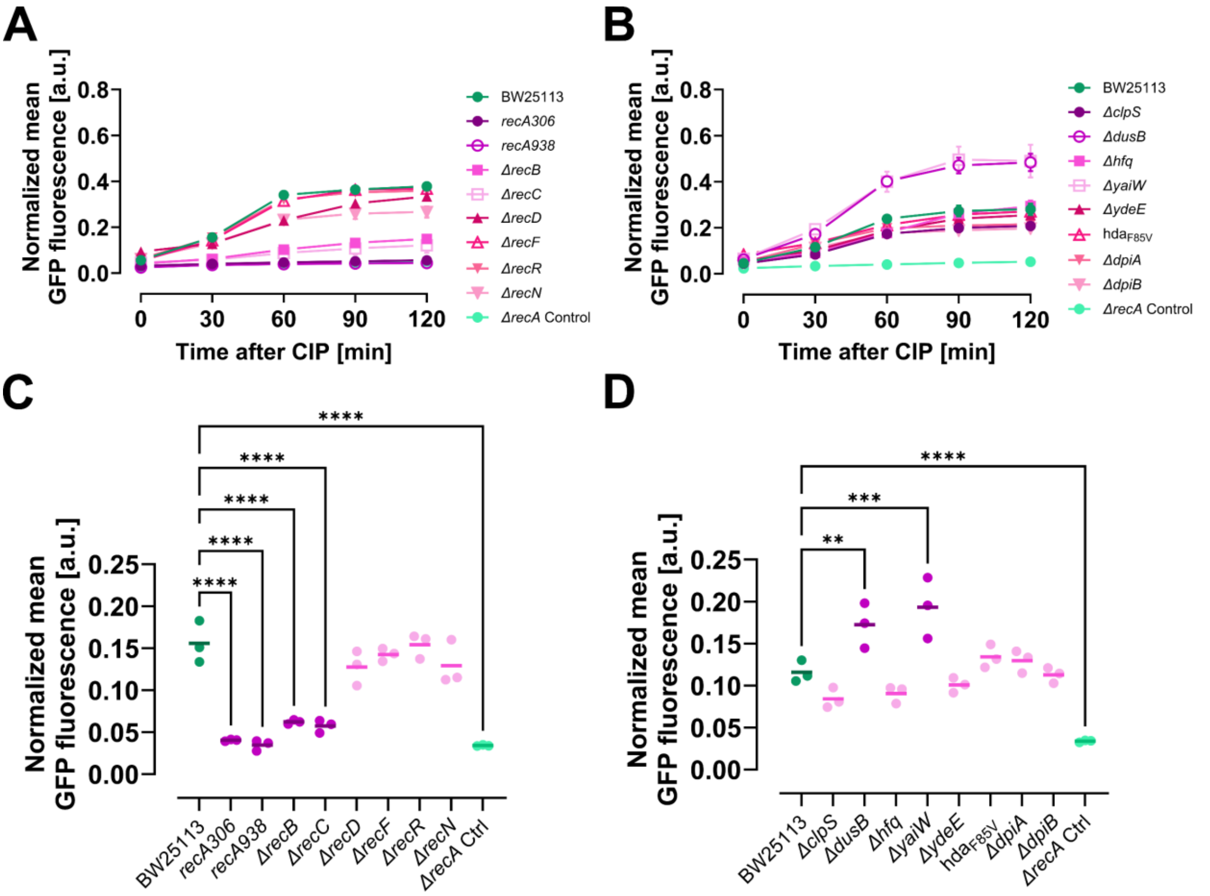


**Figure S3.** Supplementary quantifications of SOS response activity in hit strains after exposure to 10 μg/mL CIP. GFP fluorescence, regulated by the *lexA* promoter and expressed from a reporter plasmid, indicates SOS response activity and was measured in 20,000 cells per sample using a flow cytometer. (**A** and **B**) show activity measured at 30-minute intervals from just before CIP exposure until 120 minutes after., while (**C** and **D**) present only the activity after 30 minutes of CIP exposure. The assay was performed separately for strains with mutations or deletions of (**A** and **C**) genes linked to recombinational repair and the SOS response, and (**B** and **D**) genes not previously associated with DNA compaction or repair. A *ΔrecA* strain from the Keio collection (JW2669) served as negative control. SOS response activity was quantified by normalizing mean GFP fluorescence to mean forward scatter area (mean cell size). (A and B) Symbols with connecting lines represent means from three biological replicates; error bars indicate SEM. (C and D) Lines represent means from three biological replicates; dots indicate individual replicate means. The wild type (BW25113) is presented in dark green, *ΔrecA* in light green. Strains with significantly different activity from the wild type of the corresponding set of experiments are in dark magenta; others are in light magenta. Ordinary one-way ANOVA with Dunnett correction was used for comparisons; only significant differences are annotated. $^{**}P \le 0.01$; $^{***}P \le 0.001$; $^{****}P \le 0.0001$.

**Table S7.** Estimates for fixed effects from a linear mixed-effects model assessing time-dependent survival after exposure to 10 µg/mL CIP for hit strains with gene deletions linked to recombinational repair and the SOS response. Relative survival values (Fig. 9A) were $log_{10}$-transformed and modelled with exposure time (minutes) and strain (wild type as reference) as fixed effects, and biological replicates as random intercepts. The (Intercept) estimate represents the $log_{10}$ relative survival of the wild type prior to CIP exposure. The estimate for CIP exposure time indicates the change in $log_{10}$ relative survival per minute of exposure, while estimates for each deletion strain reflect their difference in $log_{10}$ relative survival from the wild type. A change of -1 in $log_{10}$ relative survival corresponds to a 10-fold (90%) reduction in survival. 95% confidence intervals are shown for each estimate, along with *P*-values derived from testing whether the fixed effects differ from zero—i.e. whether survival changes with exposure time and whether deletion strain sensitivities differ from the wild type.

| **Fixed effect** | **Estimate** | **95% Confidence interval** | ***P*-value** |
|---|---:|---|---:|
| (Intercept) | 1.102 | [0.954, 1.25] | <0.0001 |
| CIP exposure time | -0.018 | [-0.022, -0.015] | <0.0001 |
| *ΔrecB* | -1.067 | [-1.289, -0.846] | <0.0001 |
| *ΔrecC* | -1.219 | [-1.441, -0.998] | <0.0001 |
| *ΔrecD* | 0.742 | [0.52, 0.963] | <0.0001 |
| *ΔrecF* | 1.013 | [0.808, 1.218] | <0.0001 |
| *ΔrecR* | -0.098 | [-0.302, 0.107] | 0.3252 |

**Table S8.** Estimates for fixed effects from a linear mixed-effects model assessing time-dependent survival after exposure to 10 µg/mL CIP for hit strains with novel gene deletions not previously associated with DNA compaction or repair. Relative survival values (Fig. 9B) were $log_{10}$-transformed and modelled with exposure time (minutes) and strain (wild type as reference) as fixed effects, and biological replicates as random intercepts. The (Intercept) estimate represents the $log_{10}$ relative survival of the wild type prior to CIP exposure. The estimate for CIP exposure time indicates the change in $log_{10}$ relative survival per minute of exposure, while estimates for each deletion strain reflect their difference in $log_{10}$ relative survival from the wild type. A change of -1 in $log_{10}$ relative survival corresponds to a 10-fold (90%) reduction in survival. 95% confidence intervals are shown for each estimate, along with *P*-values derived from testing whether the fixed effects differ from zero—i.e. whether survival changes with exposure time and whether deletion strain sensitivities differ from the wild type.

| **Fixed effect** | **Estimate** | **95% Confidence interval** | ***P* value** |
|---|---:|---|---:|
| (Intercept) | 1.493 | [1.044, 1.942] | <0.0001 |
| CIP exposure time | -0.022 | [-0.025, -0.019] | <0.0001 |
| *ΔclpS* | -0.486 | [-1.11, 0.139] | 0.1158 |
| *ΔdusB* | -0.559 | [-1.183, 0.066] | 0.0749 |
| *Δhfq* | 0.214 | [-0.41, 0.838] | 0.4696 |
| *ΔyaiW* | -0.501 | [-1.125, 0.123] | 0.1058 |
| *ΔydeE* | 0.429 | [-0.195, 1.053] | 0.1599 |

**Table S9.** Estimates for fixed effects from a linear mixed-effects model assessing dose-dependent survival of the *ΔrecN* strain versus wild type for a range of UV doses. Relative survival values (Fig. 11C) were $log_{10}$-transformed and modelled with UV dose (J/m$^2$) and strain (wild type as reference) as fixed effects, and biological replicates as random intercepts. The (Intercept) estimate represents the $log_{10}$ relative survival of the wild type prior to UV irradiation. The estimate for UV dose indicates the change in $log_{10}$ relative survival per J/m$^2$ UV irradiation, while the estimate for *ΔrecN* reflect its difference in $log_{10}$ relative survival from the wild type. A change of -1 in $log_{10}$ relative survival corresponds to a 10-fold (90%) reduction in survival. 95% confidence intervals are shown for each estimate, along with *P*-values derived from testing whether the fixed effects differ from zero—i.e. whether survival changes with UV dose and whether the *ΔrecN* strain's sensitivity differs from the wild type.

| **Fixed effect** | **Estimate** | **95% Confidence interval** | ***P* value** |
|---|---:|---|---:|
| (Intercept) | 2.976 | [2.501, 3.451] | <0.0001 |
| UV dose | -0.077 | [-0.082, -0.071] | <0.0001 |
| *ΔrecN* | -0.604 | [-1.244, 0.036] | 0.0603 |

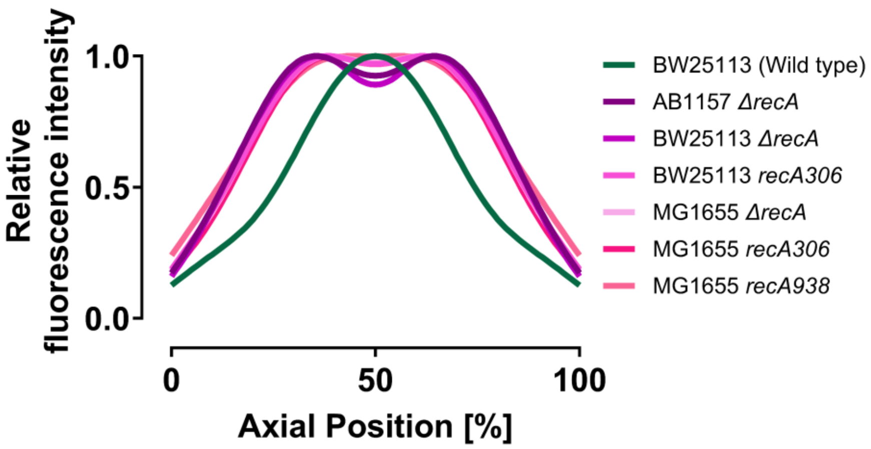


**Figure S4.** Comparison of DNA profiles along the cells' long axis after 20 minutes of CIP exposure (10 µg/mL) for various strains with *recA* deletions against the BW25113 wild type, used to evaluate compaction impairment in *recA* deletion strains.
