## Supplementary Material for "Exploring the genetic landscape of ciprofloxacin-induced DNA supercompaction in *Escherichia coli*"

#### Overview of CellProfiler measurements used for model training

To support the training of our machine learning models, we used CellProfiler (version 4, (1)) to perform a range of measurements on segmented cells from the training images. These measurements of cell and DNA features were conducted at both the image and object levels, providing a robust foundation for distinguishing DNA supercompaction phenotypes.

At the image level, CellProfiler quantified features such as area and intensity within segmented cells and DNA foci, measuring the total area occupied and perimeter length of these objects, as well as total, mean, and median intensity values. These metrics capture the broad characteristics present across images, summarizing the overall cellular composition.

At the object level, we measured a range of features for individual segmented cells and DNA foci. These features included size and shape measurements such as object area, eccentricity, and major and minor axis lengths, along with intensity metrics like integrated, mean, maximum, and minimum intensity values. More complex features included radial intensity distribution within cells, centered on DNA foci located in each cell, and intensity center shift—quantifying deviations between geometric centers and intensity gravity centers. To characterize complex shape and intensity patterns, CellProfiler quantified Zernike moments using Zernike polynomial analysis (2, 3). These moments provide a geometrical framework for identifying subtle differences in shapes and intensity distributions, aiding in classifying DNA supercompaction phenotypes. Finally, DNA foci were linked to parent cell objects, allowing for counting and calculation of mean values for foci within each cell.

All CellProfiler pipelines used in this study are available at our Zenodo repository, where further details on the specific measurements can be explored. Comprehensive descriptions for each type of CellProfiler measurement are also available in the CellProfiler online manual.

#### Training details for machine learning models

In this study, we trained two machine learning models to classify strains from the screening: the Single-Cell model and the Phenotype model. Using the graphical training interface in CellProfiler Analyst (version 3, (4)), both models were trained with a 15-parameter FastGentleBoosting classification strategy to ensure reproducibility and transparency. As our goal was to classify strains based on their similarity to different control sample phenotypes, we only included images from control samples in training datasets. These samples comprised a wild-type strain (BW25113) and a *ΔrecN* strain (JW5416), either exposed to CIP for 15-20 minutes or left unchallenged before fixation and DNA staining, representing the key phenotypic outcomes.

For training the Single-Cell model, we compiled a dataset of images from fourteen samples of CIP-exposed or unchallenged wild-type strains. From these images, we randomly fetched 2000 segmented objects and manually annotated them as either single cells or irrelevant objects. Training on the annotated cell objects resulted in a model facilitating the exclusion of irrelevant objects from further analyses. The Single-Cell model's parameters decided from the training are presented in Supplementary Table S2.

The Phenotype model was trained using a dataset of images drawn from a subset of locations across 83 of the 175 control samples from screening replicates. From this dataset, we randomly fetched 2000 cells that had been filtered by the Single-Cell model. These cells were manually annotated according to their DNA supercompaction phenotype into one of three classes: (1) wild-type phenotype displaying midcell compaction, (2) *ΔrecN* phenotype displaying quarter-position compaction, and (3) unchallenged phenotype showing multifocal distribution (Fig. 2A). Training on these annotated cells resulted in a model enabling efficient classification of each cell's DNA supercompaction phenotype. The parameters of the Phenotype model decided from the training are detailed in Supplementary Table S3.

**Testing of the trained machine learning models**

We evaluated the performance of the Single-Cell and Phenotype models using testing datasets of objects independent of those used during training. Testing involved random sampling of objects predicted by the models to belong to each class, followed by manual annotation to determine the true class of each object. These results enabled us to calculate the $F_1$ score for each class—a comprehensive performance metric. The $F_1$ score is the harmonic mean of two critical metrics: precision and recall. Precision measures the proportion of true positives among all instances predicted as positive, while recall assesses the proportion of true positive predictions within all actual positive instances. By combining these metrics, the $F_1$ score allows us to evaluate the model's accuracy in correctly identifying objects while minimizing false predictions.

To test the Single-Cell model, we randomly sampled 200 objects predicted as single cells and 200 objects predicted as irrelevant, with all objects independent from the training set. After manual annotation, we found that 153 of the predicted single cells were true positives, while 47 were false positives (Supplementary Fig. S1A). Meanwhile, 186 objects predicted as irrelevant were true negatives, with 14 false negatives (Supplementary Fig. S1A). This evaluation resulted in $F_1$ scores of 0.84 for single cells and 0.86 for irrelevant objects. The model's performance was deemed satisfactory, providing us with the confidence to advance to development of the compaction phenotype classification model.

To test the Phenotype model, we randomly sampled 200 single cells independent of the training set for each predicted compaction phenotype and manually annotated them to determine their true classifications. For cells predicted to exhibit the wild-type phenotype, 159 were confirmed as true positives, while 41 were false positives (Fig. 2B). For *ΔrecN* phenotype predictions, 122 cells were correctly identified, whereas 78 were incorrectly labelled (Fig. 2B). Meanwhile, unchallenged phenotype predictions yielded 133 true positives and 67 false positives (Fig. 2B), resulting in $F_1$ scores of 0.81 for wild-type, 0.64 for *ΔrecN*, and 0.63 for unchallenged phenotypes. The model's performance for phenotypes indicating impaired DNA supercompaction was moderate, as it often confused the two classes (Fig. 2B).

However, since both classes hold equal importance in our screening, we combined them for evaluation purposes: predictions were considered correct if a *ΔrecN* control sample was predicted to display either a *ΔrecN* or unchallenged phenotype, and vice versa. This approach substantially improved the model's prediction performance, achieving an $F_1$ score of 0.91 for the combined class with impaired DNA supercompaction. To further assess practical applicability, we used the model to predict phenotypes for all 83 control samples used to create the training dataset, calculating enrichment scores (see description below) with a lower limit of a 3:1 odds for enrichment of the phenotype. The classification was deemed correct if the enrichment score exceeded this threshold for the sample's true compaction phenotype, not differentiating between the *ΔrecN* and unchallenged phenotypes. The model provided predictions leading to 63 correct classifications, with only two samples (2.4%) misclassified and 18 (21.7%) unclassified, demonstrating reliable performance with minimal false positives. This performance was satisfactory for implementing both models into the CellProfiler pipeline used to analyze the screening images.

**Feature importance calculation**

To assess how various parameters of the Single-Cell and Phenotype models contribute to class differentiation, we quantified the feature importance for each class (Fig. 2C and Supplementary Fig. S1B). This quantification revealed which parameters were most important in distinguishing the defined classes. Model parameters dictate the weighting based on whether a feature measurement exceeds a defined threshold (positive) or falls below it (negative). For each class, we calculated a parameters' importance as the absolute difference in weighting between the positive and negative outcomes, and then normalized these values to the total importance sum across all parameters. Feature importance was determined by summarizing the cumulative importance of parameters employing the same features but utilizing different thresholds. This approach allowed us to evaluate the broader significance of each feature across various parameters, enabling a deeper understanding of how classes are differentiated. Feature importance values were calculated using a Python script available in our Zenodo repository (see Data Availability).

**Calculation of enrichment scores**

We calculated enrichment scores to assess compaction phenotypes across different samples and identify candidate strains from the screening. These scores were calculated using a Python script available in our Zenodo repository (see Data Availability) with methods adapted from the open-source CellProfiler Analyst code (4, 5). The Phenotype model first predicts a compaction phenotype for each individual cell, and the enrichment score calculation then aggregates these predictions across all cells within a sample. The script counts the cells assigned to each phenotype and models these data using a beta-binomial distribution. The enrichment score for a phenotype within a sample is calculated using the log-odds (base 10) of the likelihood that the sample contains a greater proportion of cells exhibiting this phenotype than the overall population of samples. This score therefore quantifies the odds that a sample is more enriched for a specific phenotype than what is common in the population. For instance, a sample with an enrichment score of 1 has a 10:1 odds of being enriched for the given phenotype. This method effectively ranks samples by considering both individual sample data and the collective phenotype distribution across all samples, providing a comprehensive metric for evaluating compaction phenotypes and selecting candidate strains.

### Selection criteria for candidates and poorly growing strains for re-imaging

Our selection process aimed to identify strains enriched for either the *ΔrecN* or the unchallenged compaction phenotypes, emphasizing deviations from the wild-type DNA supercompaction. We established distinct criteria based on enrichment scores to evaluate potential hits from the screening. To classify samples as enriched for the *ΔrecN* phenotype, we required enrichment scores greater than 0.477 (equivalent to 3:1 odds). A stricter criterion was applied for the unchallenged phenotype, where enrichment scores needed to exceed 1 (10:1 odds) due to generally higher scores for this class. Each replicate plate from the screening was assessed independently. When the standard criteria for the unchallenged phenotype yielded over 50 samples from a replicate plate, we increased the threshold to an enrichment score above 2 (100:1 odds), ensuring at least 30 samples remained.

Samples meeting these selection criteria underwent a quick manual image review to confirm model predictions. Samples with overcrowded cell populations in the images were often misclassified and were therefore excluded from consideration as hits. For the remaining strains, images from all replicate samples were assessed, and strains were kept on the candidate list only if none of the replicates displayed a wild-type DNA supercompaction phenotype. This selection process resulted in a total of 156 candidate strains for further investigation (Supplementary Table S4).

Certain strains included in the screening possess mutations that impair growth (6–8), which could not be accounted for under the high-throughput conditions of the screening. Consequently, these strains exhibited insufficient growth, necessitating individual culturing in tubes to ensure they reached exponential growth before assessing their compaction phenotypes. To manage the number of strains and effectively select them for re-imaging, we established specific criteria based on cell counts and enrichment scores. These criteria aimed to identify strains whose true compaction phenotypes could not be accurately assessed from the screening due to poor growth. Strains with fewer than 100 cells on average across replicates were selected for re-imaging if at least one replicate was enriched for the *ΔrecN* or unchallenged phenotypes and none were enriched for the wild-type phenotype. Additionally, strains averaging 100 to 500 cells across replicates were selected for re-imaging if they demonstrated enrichment in one replicate for the *ΔrecN* phenotype or in all three replicates for the unchallenged phenotype, provided none were enriched for the wild-type phenotype. Using this approach, we identified 93 poorly growing strains (Supplementary Table S5) that would be individually cultured for re-imaging to ensure accurate assessment of their compaction phenotypes.
